# Automated docking refinement and virtual compound screening with absolute binding free energy calculations

**DOI:** 10.1101/2020.04.15.043240

**Authors:** Germano Heinzelmann, Michael K. Gilson

## Abstract

Absolute binding free energy calculations with explicit solvent molecular simulations can provide estimates of protein-ligand affinities, and thus reduce the time and costs needed to find new drug candidates. However, these calculations can be complex to implement and perform. Here, we introduce the software BAT.py, a Python tool that invokes the AMBER simulation package to fully automate the calculation of binding free energies for a protein with a series of ligands. We report encouraging initial test applications of this software both to re-rank docked poses and to estimate overall binding free energies. We also show that it is practical to carry out these calculations cheaply by using graphical processing units in common machines that can be built for this purpose. The combination of automation and low cost allows this procedure to be applied in a relatively high-throughput mode, and thus enables new applications in early-stage drug discovery.

## 1 Introduction

Protein-ligand binding free energy calculations based on atomistic molecular simulations promise to play a growing role in drug discovery, as they provide estimates of the binding affinities of compounds proposed as drug candidates for a protein target, and thus may reduce the time and cost required for trial-and-error experimentation.^1,2^ Thus, in settings where the calculations are sufficiently fast and accurate, ^3^ one may anticipate significant savings of time and cost in early stages of drug discovery.^4–7^ It is informative to divide this broad class of methods into two subtypes: relative binding free energy (RBFE) calculations; ^8–14^ and absolute binding free energy (ABFE) calculations. ^15–23^ The former (RBFE) estimate the difference in binding free energy between two compounds by computing the change in free energy associated with a non-physical transformation of one compound to the other, in the binding site and in bulk solvent.^8,24^ Because it is easiest to carry out such alchemical transformations between compounds that are similar to each other and that adopt similar bound poses, RBFE calculations are often regarded as particularly suitable for the lead-optimization stage of drug discovery, where small chemical modifications of the initial lead compound must be selected. Interestingly, though, a recent study reports greater impact on the earlier hit-to-lead stage.^3^

In contrast, ABFE calculations estimate the standard free energy of binding of a single compound for a protein by considering a process in which a single compound is removed from the binding site into bulk solvent. This can be accomplished by a non-physical (i.e. alchemical) decoupling pathway,^15,16,25^ in which the ligand is, in effect, decoupled from the binding site and recoupled with bulk solvent; or by modeling the physical process of moving the fully coupled ligand out of the binding site^18,26–28^ into solvent. It is worth noting that any valid ABFE method must account for the free energy of releasing the ligand to bulk solvent at standard concentration.^16^ The RBFE and ABFE approaches have shown similar accuracy, relative to experiment, with computed binding free energies consistently showing root-mean-square deviations from experiment close to 1 kcal/mol.^9,12,19,25,27^

Because ABFE calculations do not involve the alchemical conversion of one compound into another, they can be applied to more diverse sets of compounds, and therefore should be a better fit to the task of screening a library of diverse compounds for binding to a targeted protein.^12,20,25,27^ However, this application is complicated by the fact that typical MD simulations are too short to allow a ligand initially modeled in one suggested pose to sample a variety of different poses. Because the chemical potential of the bound state is dominated by contributions from the most stable pose(s), this means it is essential that binding free energy calculations start from the most stable poses. Docking calculations can suggest such poses as starting points for free energy simulations, but they are not fully reliable and indeed sample a different energy landscape than an explicit-solvent MD simulation. Encouragingly, several prior studies show that ABFE calculations may be used to determine the relative stability of various plausible binding modes of a given ligand.^19,20,27^ Thus, we have the possibility of a workflow in which ABFE calculations are carried out for multiple plausible poses provided by a docking program, and the results are synthesized to provide information on which poses are most stable and on the overall binding free energy of the ligand. The present study is a step toward this goal.

ABFE calculations generally come at a higher computational cost than RBFE calculations, because a greater portion of the phase space may need to be sampled to produce well-converged results. Thus, the use of ABFE calculations to enhance pose-prediction and virtual screening has become increasingly attractive with recent increases in the time scales accessible by MD simulations, particularly through the use of inexpensive Graphics Processing Units (GPUs).^29–32^ However, the widespread adoption of these methods has remained limited by the substantial amount of human effort needed to use them. Key steps include assignment of force field parameters, construction of initial system configurations, setup of conformational restraints, equilibration of the simulation windows, execution of the production runs, and analysis of the results. Important prior contributions toward automation of ABFE calculations include the CHARMM-GUI server^33,34^ and the binding free energy estimator (BFEE).^35,36^ The former is a web-based interface that helps create input files for the various stages listed above. The latter is a tcl plug-in for VMD,^37^ with a graphical interface that creates a complete ABFE workflow starting from an initial prepared and equilibrated protein-ligand complex. However, these tools do not automate virtual compound screening with ABFE calculations. In addition, there would be great value in an open-source package written in the flexible and widely used Python programming language, as this would facilitate use, replication, customization and extension of the method.

Accordingly, in order to advance the use of ABFE calculations in drug-discovery, we have developed Binding Affinity Tool (BAT.py), an open-source, end-to-end, “black-box” package for ABFE calculations. The BAT package is written in Python and includes features that make it compatible with outputs of docking calculations, enabling its use as both a pose refinement and a ligand ranking tool. It allows the computation of the binding free energy of any ligand to a chosen receptor with minimal manual intervention, starting only from the coordinates of one or more co-crystal structures or docked complexes, along with a set of user-defined input parameters. The software also allows for user selection of the water model used, spring constants for the various restraints, number of steps for each calculation, use of hydrogen mass repartitioning (HMR),^38^ etc. To speed the calculations, it takes advantage of the high computational performance of AMBER’s *pmemd*.*cuda* sofware^39,40^ on GPUs. The BAT.py package is designed to be directly applied in the early stages of drug discovery, with the aim of using computation to reduce the time and cost of experimentation. The present study illustrates these procedures for the co-crystal structure and five docked poses of a drug-like compound bound to the second bromodomain of the BRD4 protein.

## 2 Functionality and workflow of BAT

The primary input to BAT is one or more three-dimensional structures of a given ligand and protein, each with a different pose of a given ligand and, potentially, a distinct bindingsite conformation. These structures may be experimentally determined cocrystal structures and/or crystal structures of the protein with ligand poses generated by a docking algorithm. The software also requires files specifying the force field parameters to use in the simulations, and a BAT input file containing parameters such as those defining the restraints (below), the dimensions of the simulation box, and the MD timestep. A single BAT input file can be used for multiple protein-ligand input structures, including varied poses and ligands, without additional user intervention. The output of the BAT run for a given ligand and protein is a file containing the predicted binding free energy of the ligand for each input pose. The pose with the most favorable binding free energy then is predicted as the most stable pose, and the overall binding free energy can be computed from the free energies 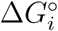 of the *N*_*pose*_ individual poses *i*:

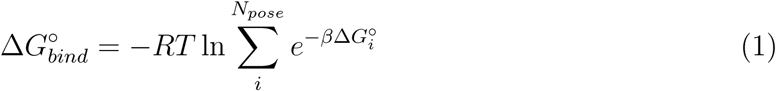

where *R* is the gas constant, *T* is absolute temperature, and *β*^−1^ = *RT*. The BAT package is written in Python and makes use of OpenBabel, VMD, MUSTANG, AMBER-Tools, and the AMBER simulation software *pmemd*.*cuda*. It is is available at the link https://github.com/GHeinzelmann/BAT.py.

The overall flow of the algorithm (Fig. 1) starts with the input structures. The first step is to assign protonation states and force field parameters to the ligand. Next, the input protein structure is aligned with a reference structure of the same protein, or a similar one, so it has the correct orientation for the various restraints to be applied. These restraints require anchor atoms on the protein and ligand, as well as three dummy atoms, which are positioned based on the input structure. Protonation states and force field parameters are then assigned to the protein, and the system is solvated with water molecules, ions needed for electroneutralit, y and, optionally, additional ions to set a desired ionic strength. Protein and ligand restraints, detailed below, are defined and imposed, and an initial equilibration simulation is carried out for each pose, while the ligand restraints are gradually released to generate starting configurations for the subsequent free energy calculations. If the ligand starting from a given pose leaves the binding site at this stage, the pose is deemed unstable, and no free energy calculation is done for it. This filter saves time by terminating calculations for ligands or poses that are clearly unstable. Additional simulations are then run to prepare the windows needed for the binding free energy calculations for each pose. The free energy calculations are then executed for all poses and the results are saved to the output file. If the input has multiple ligand poses, all are processed in the same manner.

**Figure 1:**
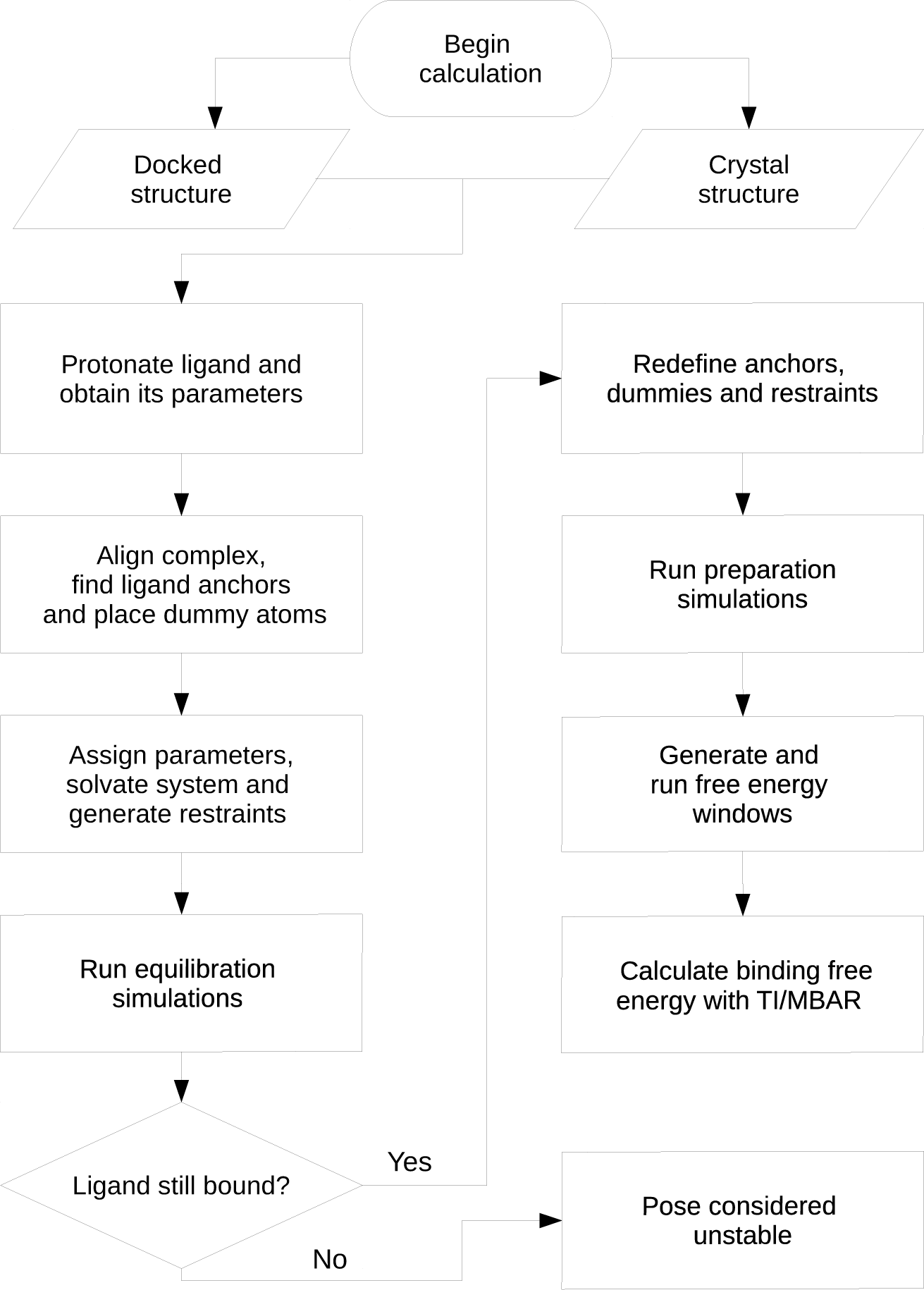
Workflow of the BAT.py software. See text for details.

## 3 Theoretical framework

This section provides the theoretical rationale for the computational procedures. For simplicity, we consider here the binding free energy of a single pose. When multiple poses are considered for a ligand, these may be combined to the overall free energy according to Eq 1.

The dissociation constant (*K*_*d*_) of a ligand-protein complex, LP, to free ligand, L, and protein, P, is related to the binding free energy by the expression: ^16^

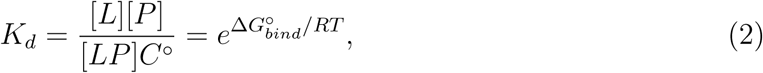

where R is the gas constant, *C*^°^ is the standard concentration of 1 M, [L], [P] and [LP] are the equilibrium concentrations of the respective species, and 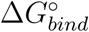 the standard binding free energy of the two molecules. In principle, the quantities [L], [P] and [LP] could be obtained from time- or ensemble-averaged ergodic simulations sampling bound and unbound states, enabling direct evaluation of *K*_*d*_ and hence 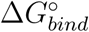. In practice, this direct approach is usually computationally intractable because of the low rate constants for the binding and unbinding processes.^1^

To overcome this limitation, one may instead obtain 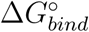 in terms of the reversible work of a dissociation process forced by artificial restraints. This dissociation process connects the bound and dissociated states with intermediate states along a pathway that may be either non-physical (“alchemical”) or physical. Either way, the overall free energy of binding may be written as a sum of terms, each corresponding to a step illustrated in Fig. 2:

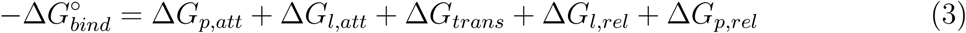

**Figure 2:**
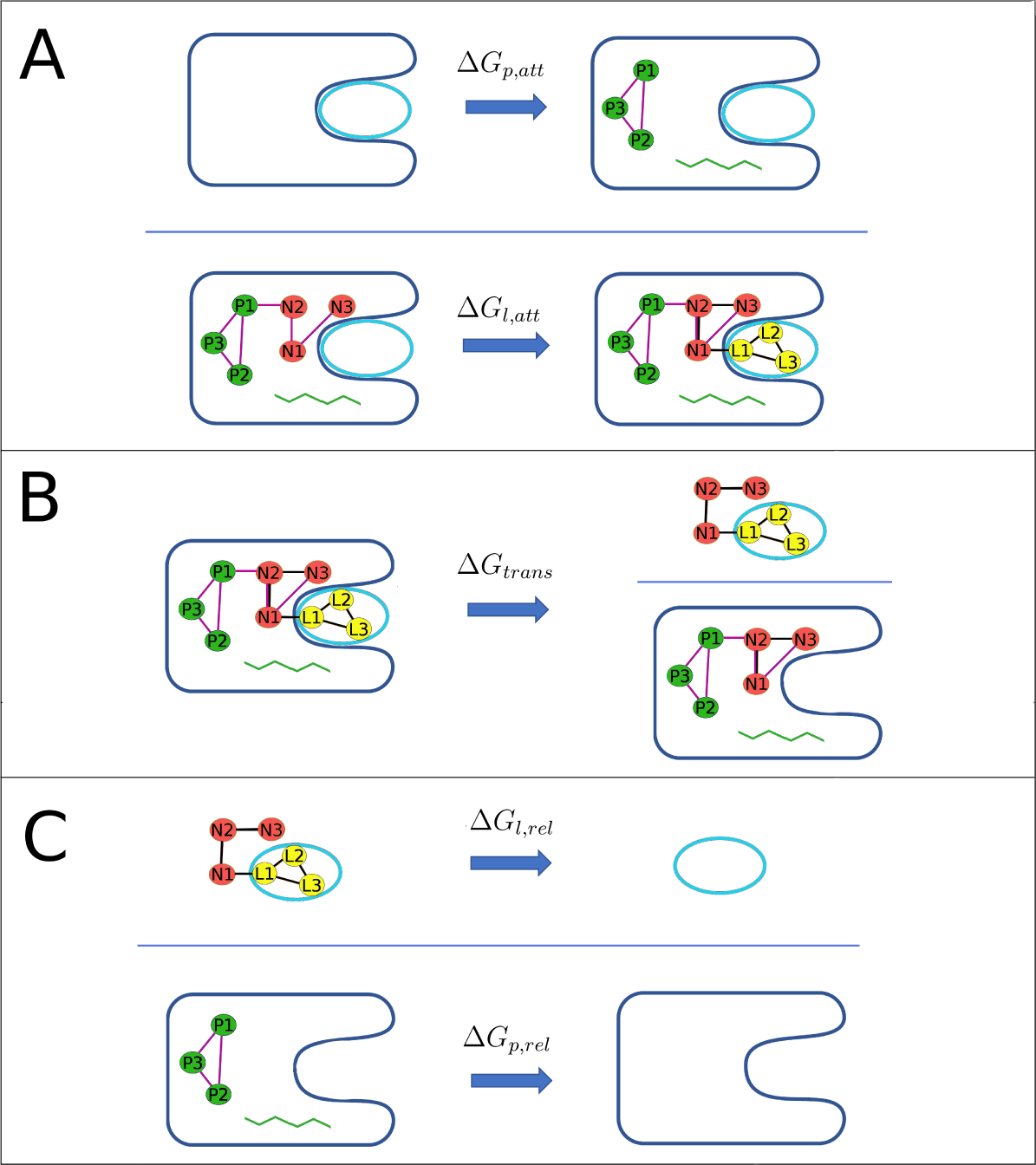
(A) Attachment of restraints, first to the protein and then the ligand. (B) Transfer ligand from binding site to bulk solvent, using the double decoupling method. (C) Release of the ligand and protein restraints in the unbound state. P1, P2 and P3 indicate protein anchor atoms; N1, N2 and N3 indicate artificial dummy atoms whose locations are fixed in the lab frame; and L1, L2 and L3 indicate ligand anchor atoms.

Above Δ*G*_*p,att*_ and Δ*G*_*l,att*_ represent the reversible work of attaching restraints, first to the protein, and then to the ligand in the context of the resulting restrained protein; Δ*G*_*trans*_ is the reversible work of transferring the restrained ligand from the restrained protein into bulk solvent; and Δ*G*_*p,rel*_ and Δ*G*_*l,rel*_ are the reversible work of then releasing the restraints that were attached in the initial steps. Note that, because the ligand and protein have negligible interactions with each other following the transfer step, the values of the two release free energies are independent of each other.

Here we use the double decoupling (DD) method,^16^ an alchemical approach which involves computing the reversible work, and hence the free energy change, of two processes: one in which the bound ligand is gradually decoupled from the protein and solvent, and a second in which the free ligand is gradually decoupled from just the solvent. The calculations use a system of artificial restraints to transfer the ligand out of the binding site into bulk solvent and facilitate conformational sampling during this process. Analogous restraints are also used to facilitate sampling during the second alchemical process, in which the ligand is decoupled from the solvent. The code accounts for the work of imposing and releasing restraints as the system progresses from one state to another. For the DD method, the transfer term Δ*G*_*trans*_ comprises two terms (Fig. 3):

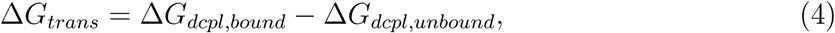

**Figure 3:**
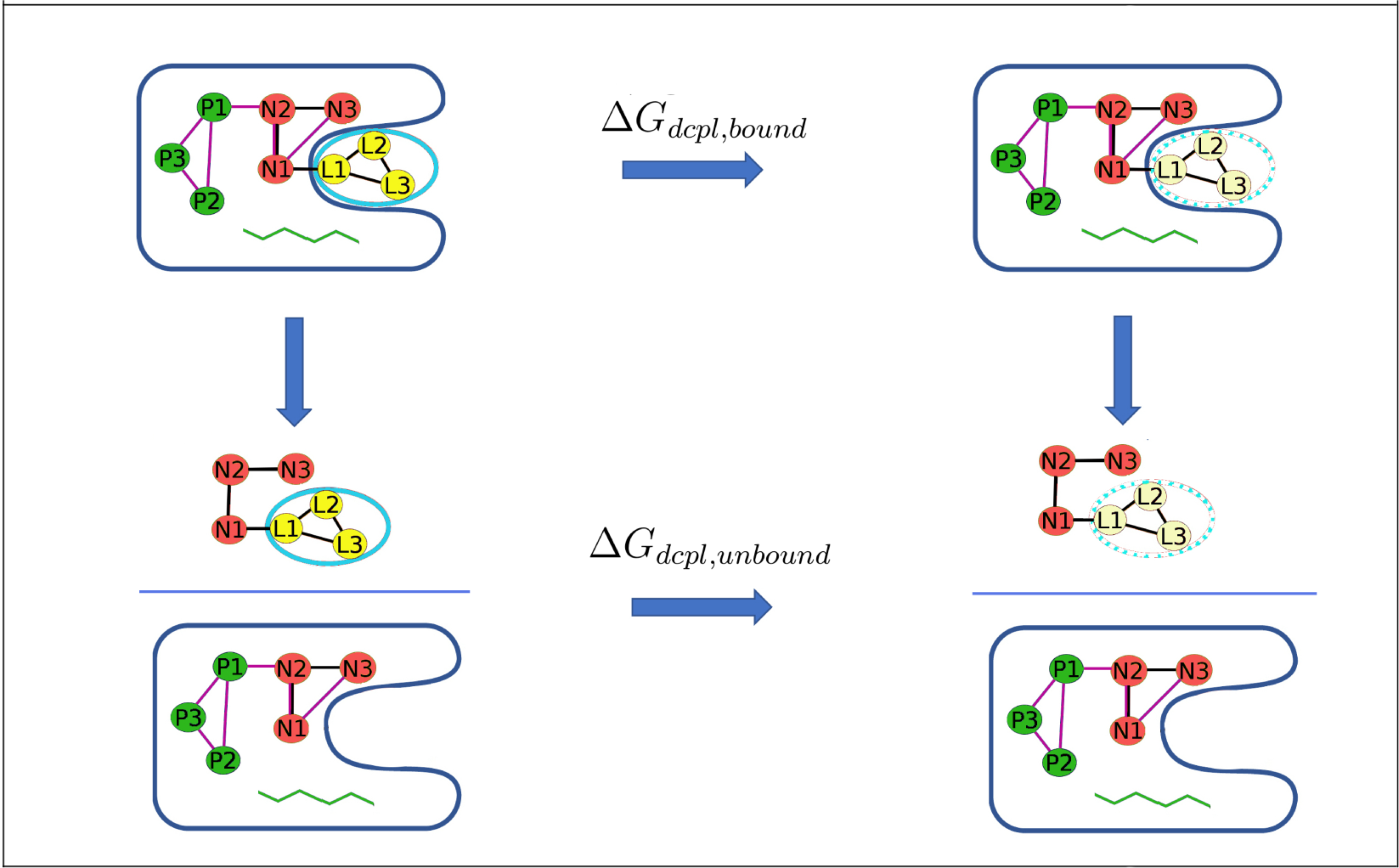
Thermodynamic cycle showing the calculation of Δ*G*_*trans*_ using the DD method, in which the restrained ligand is brought to the gas phase, both in the bound and unbound states.

Δ*G*_*dcpl,bound*_ is the reversible work of decoupling the bound ligand from the protein and solvent system to generate a ligand in the gas phase, while Δ*G*_*dcpl,unbound*_ is the reversible work of decoupling the unbound ligand from its bulk solvent environment, to generate, again, a ligand in the gas phase. Equation 4 holds because the state of the final gas-phase ligand is the same in both decoupling calculations. In the present calculations, we not only decouple the ligand from the bound and free states, but also turn off (“annihilate”) all intraligand electrostatic interactions during the decoupling process. This is because unshielded, gas-phase, electrostatic attractions can be extremely strong and cause serious conformational sampling problems, due, for example, to conformational locking by strong intramolecular hydrogen bonding.

The BAT.py code can also compute the transfer free energy Δ*G*_*trans*_ through a physical path-way, using the APR method, as done in Ref.^27^ In contrast to DD, APR needs a low-barrier physical path along which the ligand can be pulled from the binding site to bulk solvent. This is straightforward for some ligands in surface binding-sites but is difficult to assure in all cases, especially for buried binding pockets. Nonetheless, the APR implementation can still be of value, so the Supporting Information (SI) describes the present implementation and provides an illutrative sample calculation.

## 4 Methods

### 4.1 Restraints and their corresponding free energies of attachment and release

The BAT code employs essentially the same set of restraints as previously described for the APR method.^27^ Both the protein and the ligand are subject to two types of restraint. One type restrains the position and orientation of the molecule in the frame of reference of the simulation box. These translational and rotational (TR) restraints are constructed with added length, angle, and dihedral potential energy terms defined between atoms of the molecule (protein or ligand) and three dummy atoms, termed N1, N2 and N3, whose locations are fixed in the lab frame (Figure 2). These keep the protein and ligand restrained relative to the simulation box, and thus to each other.^27^ The other type of restraint is applied to internal degrees of freedom of the protein and ligand, so these limit conformational freedom. They are designed to reduce fluctuations during the calculation of the transfer free energy, and thus help convergence. This benefit must be weighed against the added computational cost of computing the attach and release free energies (Eq. 3). Note that the final free energy of binding should be the same, aside from numerical error, with or without the use of conformational restraints, and their use is, at least in principle, optional.

#### 4.1.1 Attachment and release of protein restraints

The TR restraints on the protein comprise harmonic potentials applied to one distance (D2), two angles (A3, A4), and three dihedrals (T4, T5, and T6), between the dummy atoms and three protein atoms (P1, P2, P3), which are termed the protein anchors.^27^ These protein TR restraints are present and active during the equilibration phase and throughout most of the free energy calculation. They do not affect the protein’s conformational distribution, so there is no need to compute the free energy of attaching and releasing them. The force constant for the restraint on D2 is set in the BAT.py input file with the *rec_distance_force* variable, and the force constants for A3, A4, T4, T5, and T6, are set with the *rec_angle_force* parameter. The reference values of these restraints are taken from the starting conformation. (See user manual in the SI for details.)

The conformational restraints on the protein comprise three harmonic distance restraints among the three protein anchors (P1, P2 and P3), to reduce TR fluctuations and coupling between TR motions and conformational changes. In addition, harmonic restraints may be applied to the backbone *ϕ* and *ψ* angles in a user-selected range of protein residues (Fig. 4), in order to keep this section relatively rigid when transferring the ligand from binding site to bulk, particularly when using the APR method. Backbone *ω* angles are not restrained, as these are already rather rigid, due to the double-bond character of the peptide bond. This option is activated in the BAT.py input file using the *rec_bb* variable, with the chosen residue range defined by the *bb_start* and *bb_end* parameters. The spring constants for the protein distance and backbone dihedral conformational restraints are specified with the *rec_discf_force* and *rec_dihcf_force* variables, respectively.

**Figure 4:**
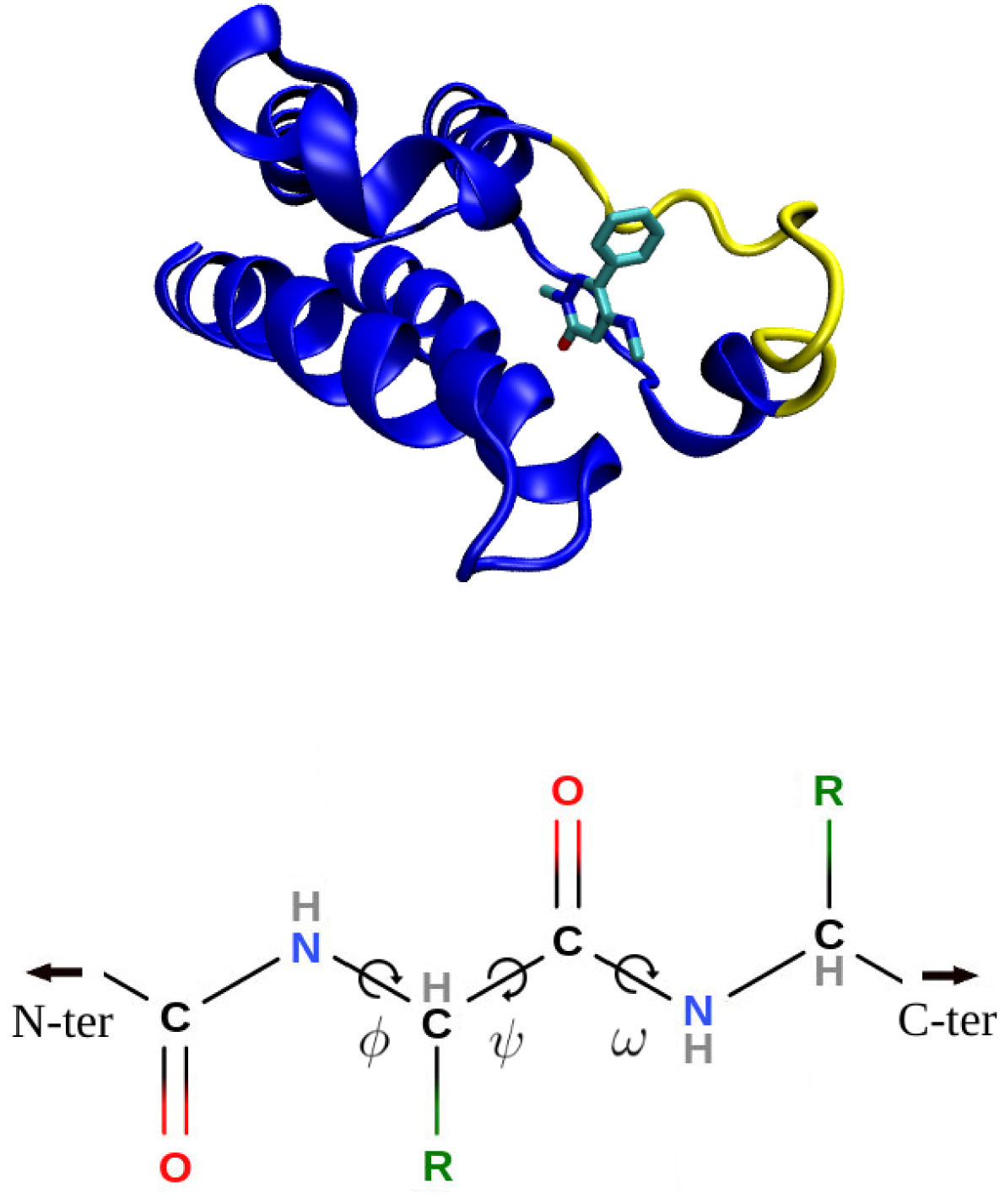
(*top*) Second BRD4 bromodomain with the restrained backbone section (part of the ZA-loop) in yellow, the rest of the protein in blue, and the ligand colored by element. The structure is from the 5uf0 cocrystal structure. (*bottom*) Protein backbone showing Ramachandran torsion angles. The optional backbone restraints in BAT.py are applied to only *ϕ* and *ψ* angles.

The free energies of attaching, (Δ*G*_*p,att*_), and releasing (Δ*G*_*p,rel*_) the protein conformational restraints are calculated in the absence of the protein TR restraints (top and bottom processes in Fig 2), using a number of simulation windows with *N*_*w*_ values of the restraint spring constants, between zero and its full value, corresponding to *N*_*w*_ windows. At each window, *i*, the spring constant for a given restraint, *r*, is defined by:

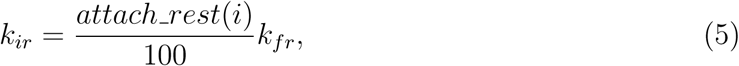

where *k*_*ir*_ is the spring constant, *k*_*fr*_ is its full value defined in the input file, and *attach_rest(i)* is the multiplying factor associated with each window. These factors are defined by the *attach_rest* array in the BAT input file, and should go from 0 to 100. The same factors are used for all of the attach and release calculations for the protein and the ligand. Note, however, that the free energy of the ligand TR release is computed semi-analytically rather than with simulations. The free energies are obtained from the trajectories of all windows, using the multistate Bennett acceptance ratio (MBAR) method,^41^ according to the equation:

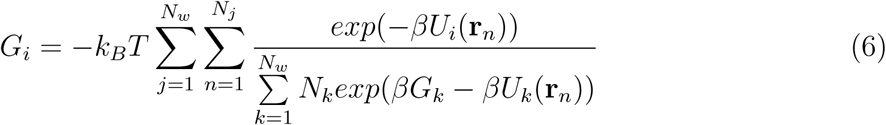

Here the subscripts *i, j*, and *k* index the simulation windows; *n* indexes the *N*_*j*_ samples from window *j*, each with coordinates **r**_*n*_, and *N*_*k*_ is, analogously, the number of samples from window *k*; *G*_*i*_ and *G*_*k*_ are the free energies of windows *i* and *k*, respectively; *β* = 1*/k*_*B*_*T*; *U*_*i*_(**r**_*n*_) is the potential energy from the restraints defined in window *i* acting on the coordinates **r**_*n*_, which correspond to the *n*th sample from window *j*. Thus, *U*_*i*_(**r**_*n*_) is given by

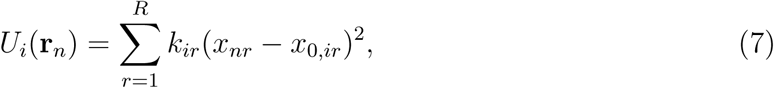

where *R* is the number of restraints being attached or released; *x*_*nr*_ is the value in sample (or frame) *n* from window *j* of the internal coordinate (distance, angle, or torsion) corresponding to restraint *r*; and *k*_*ir*_ and *x*_0,*ir*_ are, respectively, the spring constant and equilibrium value for the *r*th harmonic restraint in window *i*. The program Pymbar^41^ is invoked by BAT to solve Eq. 6 self-consistently for the free energies across the windows. The free energy difference between the initial (unrestrained) and final (fully restrained) states is then obtained directly from this set.

#### 4.1.2 Attachment and release of ligand restraints

Like the protein, the ligand is subject to harmonic TR and conformational restraints.^27^ The TR restraints again comprise a distance, D1, two angles, A1 and A2, and three torsions, T1, T2, T3, defined relative to the three fixed dummy atoms N1, N2 and N3. The spring constant for D1 is set in the BAT.py input file using the *lig_distance_ force* variable. For the angle and dihedral TR restraints, the spring constant is defined by the *lig_angle_force* parameter. The reference values of these restraints are taken from the initial coordinates. The ligand conformational restraints include harmonic potentials on the three distances between its anchor atoms L1, L2 and L3 (Fig. 2). In addition, essentially all dihedral angles are also restrained to make each ligand rigid and thereby accelerate convergence. For simplicity, torsions within rings are not excepted from the set of restraints, although they are not always necessary. The BAT.py script automatically assigns these restraints for each ligand. It uses the ligand’s AMBER parameter/topology (prmtop) file (Fig. 5) to identify all proper dihedral terms not involving a hydrogen atom, and assigns a restraint to one arbitrarily chosen dihedral term for each central bond. The spring constants for the ligand’s internal distance and dihedral restraints are set in the BAT.py input file via the *lig_discf_force* and *lig_dihcf_force* parameters, respectively, and their reference values are taken from the starting coordinates. Fig. 5 illustrates the assignment of 14 conformational dihedral restraints for the ligand from cocrystal structure 5uf0.

**Figure 5:**
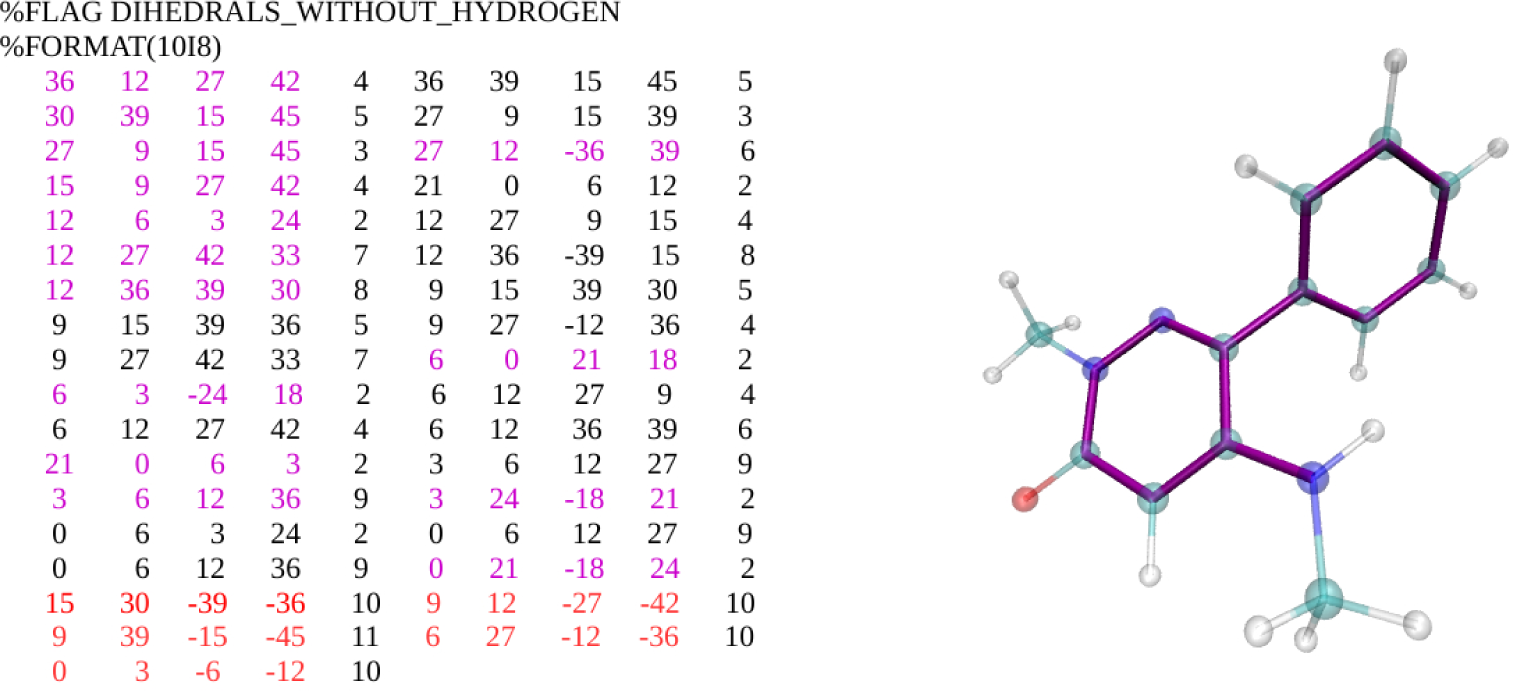
Example of ligand dihedral restraints, for PDB ID 5uf0. Left: Section of AMBER parameter/topology file listing all the ligand dihedrals that do not include a hydrogen atom. Each row lists two torsions in terms of five indices; the first four map to specific atoms and the fifth maps to the associated force field parameters. Dihedrals restrained in the BAT procedure are highlighted in purple font, with redundant ones in black and improper dihedrals in red. Right: Ligand from 5uf0 with restrained torsions highlighted with purple bonds. Cyan: carbon. White: hydrogen. Red: oxygen. Blue: nitrogen.

The free energies of attaching and releasing the ligand restraints may be separated into conformational and TR parts:

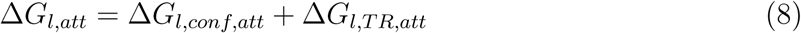

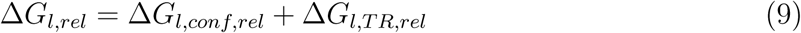

During the attachment stage (*att*), the ligand is in the binding site of the restrained protein. The conformational restraints are applied first, yielding the free energy change Δ*G*_*l,conf,att*_ for making the ligand essentially rigid. The TR restraints are then applied, yielding the free energy change (Δ*G*_*l,T R,att*_) for restraining the ligand in the binding site. During the release stage (*rel*), Δ*G*_*l,conf,rel*_ is computed with the ligand in a separate simulation box with no TR restraints present. The values of Δ*G*_*l,conf,att*_, Δ*G*_*l,T R,att*_, and Δ*G*_*l,conf,rel*_ are calculated the same way as the protein conformational restraints, using MBAR (Eqs. 5, 6 and 7), with simulation windows having intermediate values of the harmonic spring constants, also defined by the *attach rest* input array. The final term in Eq 9, Δ*G*_*l,T R,rel*_, is calculated by numerical quadrature of the following integral, which is based on Euler angles and spherical coordinates:

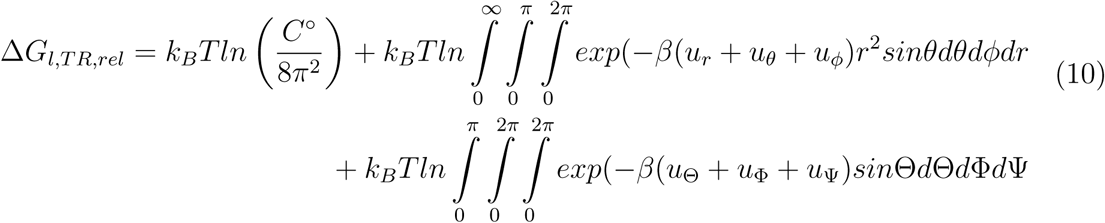

Here *C*^°^ is the standard concentration, 1 M = 1/1661Å^3^, and *r, θ* and *ϕ* are the distance D1, angle A1, and dihedral T1, respectively. ^27^ In the last term on the right, which integrates over ligand orientation, Θ is the angle A2, Φ is the dihedral T2, and Ψ is the dihedral T3. These are three Euler angles which define the orientation of the ligand in space. The harmonic potential applied to the distance *r* has the form:

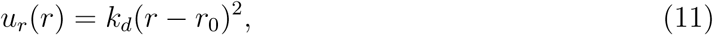

with *k*_*d*_ the *lig_distance_force* spring constant and *r*_0_ its reference value. A similar expression is used for restrained angles and dihedrals:

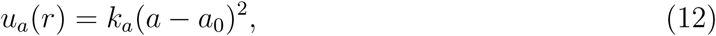

with *a* a given angle/dihedral, *k*_*a*_ the *lig_angle_force* spring constant, and *a*_0_ its reference value. The value of *r*_0_ is the reference distance, D1, from dummy atom N1 to ligand atom L1, in the bound state; this is always sets to 5.00 Å by construction (Section 4.3.1).

### 4.2 Transfer of ligand from binding site to bulk solvent

The double decoupling approach uses two non-physical paths to calculate the transfer free energy of the fully restrained ligand from binding site to bulk, Δ*G*_*trans*_ (Eq. 4 and Figure 3). Here, each of the terms on the right side of Eq. 4 is further separated into an electrostatic component and a Lennard-Jones component:

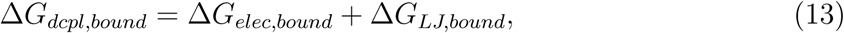

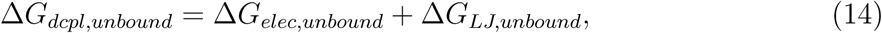

Here Δ*G*_*elec,bound*_ is the free energy change for discharging all the atomic partial charges of the bound ligand, and Δ*G*_*LJ,bound*_ is the free energy change for turning off all LJ interactions between the (electrically discharged) bound ligand and its environment. This process turns off all intraligand electrical interactions, but preserves the intra-ligand LJ interactions. The LJ decoupling term is computed with soft-core potentials as implemented in AMBER, in order to smoothly switch off the LJ interactions and thus avoid numerical problems at the transformation end-points. The free energy for ligand decoupling in bulk, Δ*G*_*dcpl,unbound*_, follows the same decoupling procedures, but for the ligand in a simulation box of solvent without the protein. Running the decoupling calculations for the ligand with its conformational restraints present avoids numerical challenges that can otherwise result from large changes in the conformational preferences of the ligand between the coupled and decoupled (gas phase) states.^42^ The BAT software allows these decoupling free energies to be computed via thermodynamic integration with Gaussian quadrature (TI-GQ) or MBAR, using the pyMBAR software, based on outputs available from the AMBER simulations. The choice of method is dictated by the *dd_type* parameter in the BAT input file.

### 4.3 Calculation definition and setup

As noted in Section 2, BAT can accept as input either a single protein-ligand cocrystal structure or a single protein structure and a set of ligand poses generated with a docking program. As detailed in the user manual, the *calc_type* parameter is used to choose between these. The processing of the input structures to generating computed binding free enegies is detailed in the following subsections.

#### 4.3.1 Anchor atoms and dummy atoms

The user identifies the desired protein anchor atoms (P1, P2 and P3; Section 4.1.1) with the *P1, P2* and *P3* variables, and provides a reference protein structure for the orientation of the target protein. It is important that the protein anchor atoms be chosen to work for the reference structure, as any additional structures of the protein will be aligned to it in the source of the free energy calculations. The alignment of the complex relative to the protein reference structure is done with the program MUSTANG,^43^ by first aligning the two protein sequences and then finding the optimal superposition of the two structures. Thus, the reference structure does not need to have the exact same sequence as the target one, so one may use only one reference for a set of similar proteins, with equivalent residues as the three protein anchors. The BAT.py procedure automatically assigns the ligand anchor atoms, L1, L2 and L3, and sets up the coordinates of the N1, N2 and N3 dummy atoms. This is done using a procedure that avoids possible gimbal-locking, which could result if the angle between three adjacent atoms of a given dihedral approached 0^°^ or 180^°^. Gimbal-locking can cause large restraint forces, leading to instabilities and crashes during the simulation, and therefore should be avoided.

The aligned protein-ligand complex is used to select the ligand anchor atoms and position the dummy atoms. First a “strike zone” is defined for use in identify the L1 ligand atom (Fig. 6, left). The strike zone is a square of side-length 2\**l1_range*, oriented perpendicular to the *z* axis. The center of this square has *x* and *y* coordinates given by *x*_*P 1*_ + *l1_x* and *y*_*P 1*_ + *l1_y*, respectively, where *x*_*P 1*_ and *y*_*P 1*_ are the *x* and *y* coordinates of atom P1 (Fig. 6). Here *l1_x, l1_y* and *l1_range* are user-defined input parameters. Guidelines for selecting these parameters are provided in the user manual. The L1 anchor atom will be the ligand atom with *x* and *y* coordinates inside the strike zone and with the lowest *z* distance from P1, with the requirement that this distance is between the user-defined parameters *l1_z* (minimum) and *l1_zm* (maximum). The minimum value ensures that the N1 dummy atom can be placed between P1 and L1 in the *z* axis, and the maximum value avoids finding L1 if the ligand has left the binding site.

**Figure 6:**
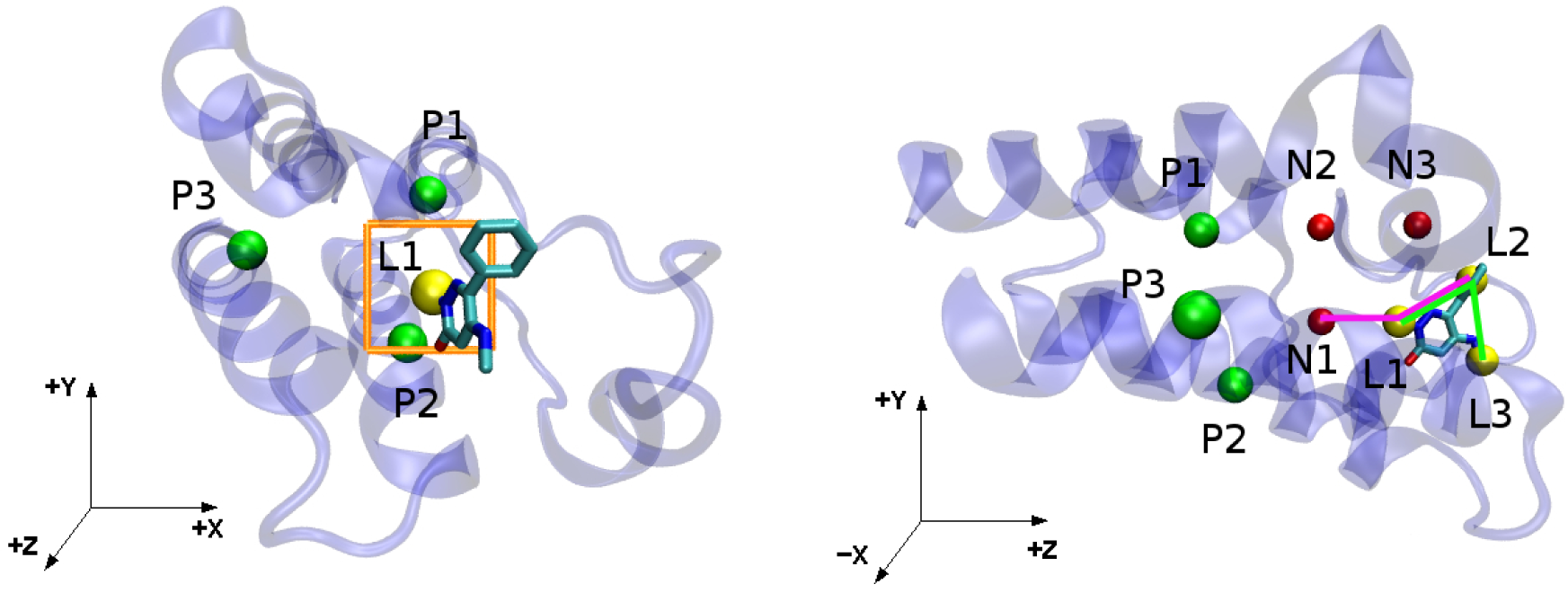
Definition of ligand anchor atoms and dummy atoms for the co-crystal structure 5uf0. Left: Strike zone (orange square), ligand anchor atom L1 (yellow), and protein anchor atoms (green). Right: Same system rotated relative to the left-hand panel, illustrating the definition of the L2 and L3 ligand anchor atoms (yellow). The N1-L1-L2 and L1-L2-L3 angles are shown with green and purple lines, respectively, and dummy atoms are shown in red.

The first dummy atom, N1, is assigned the same *x* and *y* values as L1 but with a *z* distance of 5.0 Å from L1. The whole system is now rotated around the *z* axis, so that P1, N1 and L1 have the same *x* coordinates and thus are all located in the same *yz* plane. The second dummy atom, N2, is then placed with the same *z* value as N1, but with the *x* and *y* coordinates of P1. Dummy atom N3 is then positioned with the same *x* and *y* coordinates as N2, but with a distance from N2 in the *z* axis having the same value as the magnitude of the N1-N2 distance. Now P1, L1, N1, N2 and N3 are all located in the *yz* plane, as shown in the right hand panel of Fig. 6.

The remaining ligand anchor atoms, L2 and L3, are now selected. The L2 anchor is defined as the ligand atom which provides an N1-L1-L2 angle as close as possible to 90^°^, while having an L1-L2 distance between the minimum and maximum specified in the input file as *min_adis* and *max_adis*, respectively. Setting a minimum distance between anchors L1 and L2 prevents the application of excessively large forces on dihedral restraints, which could result from a small lever arm. The maximum distance parameter is not as critical, but is meant to avoid choosing ligand anchor atoms far from the binding site. If no L2 atom can be found inside the specified distance range, as may occasionaly occur if the ligand is very small, the two parameters can be adjusted in the BAT.py input file. (If one finds this happens too often for a given application, the availability of the Python code affords a skilled user the opportunity to develop a custom version that, for example, automates the optimization of these parameters for each ligand.) The analogous protocol is then used to choose L3 based on the positions of L1 and L2, but now keeping the L1-L2-L3 angle as close as possible to 90^°^ and the L2-L3 distance within the specified distance range. The input parameters listed here only have to be defined once for a new protein system, and generally will work for any ligand that binds to the same binding pocket. This protocol ensures that the N2-N1-L1 and N1-N2-P1 angles are 90^°^, minimizing the forces applied by the T1 and T4 dihedral restraints on the P1 and L1 atoms.^27^

#### 4.3.2 Force field options, solvation, and ionization

The protonation states of the protein’s ionizable groups are predetermined by the user-selected residue templates. Protonation states of the ligand are set with the program open-babel,^44^ assuming a pH of 7.4. (This selection will be generalized in future versions.) The current BAT software automates free energy calculations using any AMBER protein force field, chosen with the *receptor_ff* variable. Ligand parameters are assigned with the AM1-BCC charge model^45^ and either version 1 or 2 of the General AMBER Force Field (GAFF),^46,47^ chosen with the *ligand_ff* input variable. Currently supported options for the water model, chosen with the *water_model* parameter, are TIP3P,^48^ TIP4PEw^49^ or SPC/E.^50^ The types of any dissolved ions are specified with the *cation* and *anion* input variables, and the ions are assigned Joung and Cheatham parameters appropriate to the selected water model. ^51^ This selection could easily be changed by modification of the Python code.

The Ambertools tleap software^39,40^ is used to solvate the protein-ligand complex and the dummy atoms in a box of water molecules. The BAT.py code allows the user to choose the number of water molecules in the box and the water padding in the *x* and *y* axes, using the *num_waters, buffer_x* and *buffer_y* parameters, respectively. The dependent variable is the water padding in the *z* direction, which is calculated using an efficient iterative relaxed Newton-Raphson approach, based on the cross-sectional *xy* area of the box, the requested number of water molecules, and the average tleap atomic density for each water model. With the *neutralize_only* keyword, dissolved counterions are added if needed for charge neutralization. With the *num_cations* parameter, a number of additional cations are also added for a desired ionic strength, with the same number of anions added for neutrality.

The binding free energy calculations also require simulations of the free protein and the free ligand. These calculations are automatically set up as follows. The size of the ligand box is set in the input file using the *lig_buffer* parameter; this defines the water padding in all three Cartesian axes. Counterions are added to neutralize the ligand, as needed. The variable *num_cat_ligbox* sets a user-defined number of additional cations, and the number of anions again is the dependent variable. The variables to create the box with the *apo* protein are the same used previously for the protein-ligand system, such as the number of waters, water model, *x* and *y* buffering, and number of cations.

### 4.4 Simulation procedures

#### 4.4.1 Energy minimization and heating

Each system prepared above is energy minimized, with the protein restraints fully turned on. Molecular dynamics is then run as the system is heated, over 100ps, from 10K to the desired simulation temperature of 298.15 K, at constant volume, and using a Langevin thermostat^52^ with a collision frequency of 1.0 ps^−1^. Then a series of brief (15 ps) simulations is run with the pressure held at 1 bar with the Monte Carlo barostat.^53^ This procedure allows the volume to adjust while avoiding possible crashes caused by excessive shrinking of the initial box. Once this step is concluded, the system is ready for all subsequent runs, which are performed at constant temperature and pressure. The simulation temperature, thermostat collision frequency and barostat type can be chosen using the *temperature, gamma_ln* and *barostat* variables, respectively.

#### 4.4.2 Equilibration and preparation

This stage prepares each initial protein-ligand complex for the free energy calculations detailed in Section 4.5. The first step is to relax the initial ligand-protein complex so that it either declares itself as unstable, and thus not worth further analysis, or else settles into a nearby free energy minimum which becomes the starting structure for the binding free energy calculation. A sequence of molecular dynamics simulations is run with weaker and weaker ligand TR restraints, but without any ligand conformational restraints, until the final simulation is performed with the ligand free in the binding pocket. The number of simulations, the scaling of the ligand TR force constants at each simulation, and the number of steps for each simulation, are defined with the variables *release_eq, eq_steps1* and *eq _steps2*. During this process, the protein TR restraints and the distance restraints among the P1, P2 and P3 anchors are maintained. If the protein backbone dihedral restraints are in use, one may either maintain them or turn them off so the protein can fully adapt to the docked ligand. This choice is controlled by the the *bb_equil* variable.

Once this relaxation is complete, an attempt is made to set up a fresh set of ligand restraints, using the procedures in Section 4.3.1. If an L1 anchor can no longer be identified inside the strike zone and within the maximum allowed P1-L1 *z* distance, *l1_zm*, the ligand is considered to have left the binding site during equilibration. The initial pose is then considered unstable, and no free energy calculation is done. However, if an L1 anchor can be identified, then the dependent anchors and dummy atoms are reassigned, all restraints are given reference values corresponding to the final equilibrated structure, and the simulation box is rebuilt, re-minimized, and re-equilibrated with all restraints in place. The number of MD steps in this second “preparation” equilibration is specified with *prep_steps1*. The resulting system structure is used to initiate calculation of the binding free energy (Section 4.5).

### 4.5 Free energy calculations

The calculated binding free energy is a sum of contributions, detailed in Sections 3 and 4; see Eqs. 3, 4, 8, 9, 10, 13, and 14. Each window of each free energy calculation is independently equilibrated, and then a production simulation is used to collect data. The number of equilibration and production MD steps for all windows of each component are set using the *[component]_steps1* and *[component]_steps2* input parameters, respectively, where *[component]* is the letter code for the free energy component (Table 1), *steps1* is the number of equilibration steps, and *steps2* is the number of production steps.

**Table 1:**
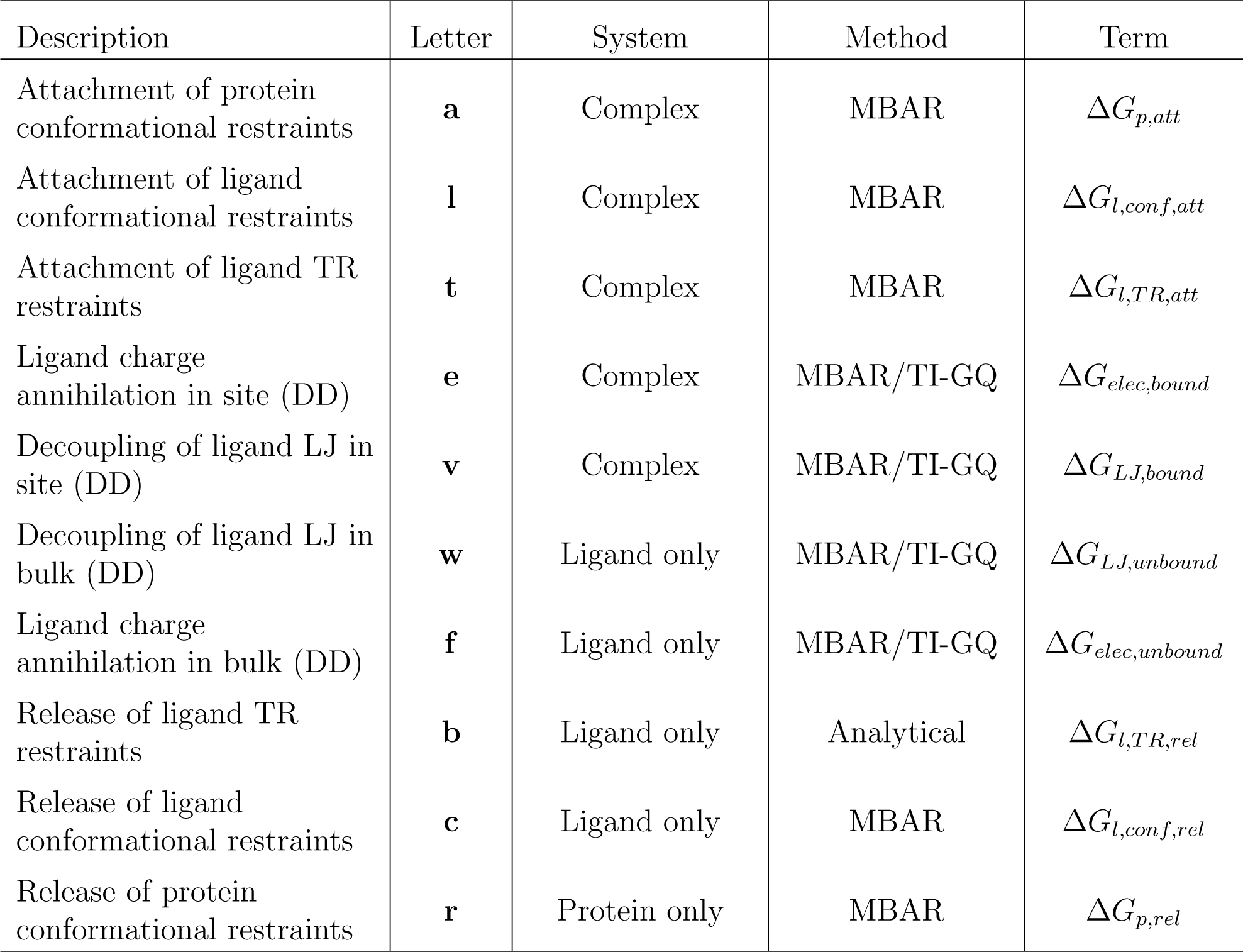
Letter codes used in specifying simulation parameters to be applied in computing the free energy contributions to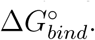 See main text and Eqs. 3, 4, 8, 9, 10, 13, and 14 for definitions of the terms. System: the molecular system simulated to compute each term. Method: the free energy method used to compute each term.

| Description | Letter | System | Method | Term |
| --- | --- | --- | --- | --- |
| Attachment of protein conformational restraints | **a** | Complex | MBAR | $\Delta G_{p,att}$ |
| Attachment of ligand conformational restraints | **l** | Complex | MBAR | $\Delta G_{l,conf,att}$ |
| Attachment of ligand TR restraints | **t** | Complex | MBAR | $\Delta G_{l,TR,att}$ |
| Ligand charge annihilation in site (DD) | **e** | Complex | MBAR/TI-GQ | $\Delta G_{elec,bound}$ |
| Decoupling of ligand LJ in site (DD) | **v** | Complex | MBAR/TI-GQ | $\Delta G_{LJ,bound}$ |
| Decoupling of ligand LJ in bulk (DD) | **w** | Ligand only | MBAR/TI-GQ | $\Delta G_{LJ,unbound}$ |
| Ligand charge annihilation in bulk (DD) | **f** | Ligand only | MBAR/TI-GQ | $\Delta G_{elec,unbound}$ |
| Release of ligand TR restraints | **b** | Ligand only | Analytical | $\Delta G_{l,TR,rel}$ |
| Release of ligand conformational restraints | **c** | Ligand only | MBAR | $\Delta G_{l,conf,rel}$ |
| Release of protein conformational restraints | **r** | Protein only | MBAR | $\Delta G_{p,rel}$ |

Following completion of the simulations, BAT.py computes each free energy component using the methods listed in Table 1; i.e., TI-GQ, MBAR, and/or analytical. This analysis uses options set in the previous stages, such as *components, dd_type, lambdas*, and *weights*. The trajectories for every window are also split into *blocks* blocks and the free energies are computed separately for each block. This feature is helpful to check for convergence, as large variations across blocks signals convergence problems during the calculations. We also report the standard deviation across blocks as a conservative estimate of the uncertainty in each free energy term:

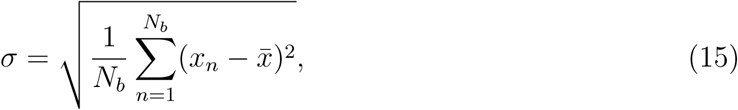

Here 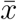 is the free energy calculated using the whole trajectory, *x*_*n*_ is the free energy calculated for each block, *N*_*b*_ is the number of blocks, and *σ* is the standard deviation. The uncertainties computed in this way for each free energy term are added in quadrature to obtain the reported uncertainty of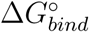.

### 4.6 Protein-ligand test system

We ran illustrative free energy calculations with BAT.py for BRD4(2), the second bromodomain of the BRD4 protein, with ligand 2-methyl-5-(methylamino)-6-phenylpyridazin-3(2H)-one (left of Fig. 7), which we will refer to as 89J, its PDB HETID. Binding free energies were computed for the cocrystal structure, 5uf0, and also for five ligand poses generated by computational docking. The docking workflow uses a modified version of the one explained in the CELPPade tutorial,^54^ which is associated with the CELPP blinded prediction challenge.^55^ It uses Autodock Vina^56^ for the docking and Chimera^57^ for protein and ligand setup and to convert the output files to pdb format. The protein structure used for docking is 5uez, ^58^ as this was identified by CELPP as having the ligand with the largest maximum common substructure (LMCSS) with our target ligand (right of Fig. 7). The docked poses were characterized by calculating their structural RMSDs relative to the reference 5uf0 crystal structure, both before and after equilibration by MD. The program VMD was used to compute the RMSD values: the two protein structures were aligned and the RMSD Calculator plug-in was used, accounting for the symmetry of the phenyl group of the target ligand.

**Figure 7:**
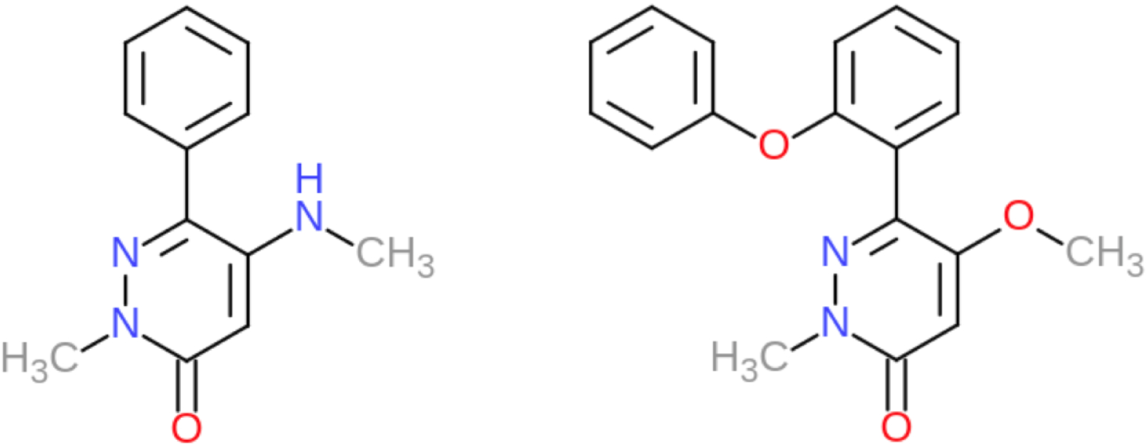
Left: Structure of the ligand, 2-methyl-5-(methylamino)-6-phenylpyridazin-3(2H)-one (89J) considered in the present free energy calculations. Right: Structure of the ligand, 5-methoxy-2-methyl-6-(2-phenoxyphenyl)pyridazin-3(2H)-one ligand, from the cocrystal structure 5uez, which provided the receptor structure for the present docking calculations.

### 4.7 Computational details

All simulations invoked by BAT.py use *pmemd*.*cuda* from AMBER18, with rectangular periodic simulation boxes. The AMBER “cut” parameter was set to 9.0 Å, and long-range electrostatics were calculated using the PME method. All bonds involving hydrogen were constrained by using the SHAKE algorithm.^59^ The three dummy particles, N1, N2, and N3, are assigned zero charge, zero Lennard-Jones radius and well-depth, and a mass of 220 Da, with their Cartesian coordinates restrained by a harmonic force constant of *k* = 50*kcal/*(*mol*. Å^2^). We used hydrogen mass repartitioning (HMR)^38^ in all simulations, and an MD with a time step of 4 fs. However, the use of HMR is optional, and is selected with the *hmr* variable. Other simulation options, such as the cutoff value, time step and output frequency, can be set in the BAT.py input file with the same variables used in the *pmemd*.*cuda* simulation input file. The input parameters for the binding free energy calculations, such as *λ* values, simulation times, force constants and water model, are included in the Supporting Information (SI).

## 5 Results and Discussion

We used BAT.py to carry out illustrative calculations of the binding free energy of BRD4(2) with its experimentally characterized ligand 89J. Calculations were done starting with the crystal structure 5uf0, and also, independently, with five distinct poses generated by using Autodock Vina to dock this ligand into the structure of BRD4(2) solved with a similar ligand (Fig. 7). Here, we report on the accuracy of the calculations and consider the potential of this type of calculation to distinguish correct from incorrect poses, with the caveat that this single test case can only serve as initial proof of principle. We then report the computational effort required for these calculations, and discuss the suitability of our software for use in a high-throughput scenario.

The calculations presented here are automated, given a set of basic input files and parameters, so they can be reproduced and generalized. The software is available on GitHub, as are a tutorial, a more complete User Guide, and the input files needed to replicate the present calculations. ^60^ Also included are the input parameters needed to run similar calculations on several other protein-ligand systems, and docking scripts that can be used to prepare the systems for BAT calculations. These scripts include the system preparation using Chimera, sample files, and a bash script to perform the docking automatically using AutoDock Vina.

### 5.1 Binding free energies and pose evaluation

The binding free energy computed with the crystallographic pose is −6.1 kcal/mol, a diference of −0.9 kcal/mol when compared to the experimental result of −5.2 kcal/mol ^58^ (Table 2 and Table S1 from the SI). In addition, binding free energies computed with the two docked poses whose RMSDs are at most 2.0 Å (poses 2 and 5) show good consistency with the crystal structure result, and are within 1.5 kcal/mol of experiment (Table 2). However, binding free energies computed with the less accurate docked poses (1, 3, and 4), with initial RMSD values ≥5.4 Å), are more positive by at least 3 kcal/mol, and hence less favorable than those obtained with the more accurate poses (Table 2). Thus, the absolute binding free energy calculations yield reasonable agreement with experiment and distinguish accurate from inaccurate binding poses, as hoped. Interestingly, during the equilibration phase of the calculations, the RMSD of the crystallographic pose rose somewhat, from 0.0 to 1.3 Å, whereas both poses 2 and 5 moved closer to the crystallographic pose. Note, however, that these values are for single conformational snapshots, and that the RMSD values fluctuate in the course of a simulation.

**Table 2:** Summary of computational results. Top two data rows give the structural root-mean-square deviations (RMSDs, Å) from the reference crystal pose in 5uf0, of the various ligand poses before (Initial) and after (Equilibrated) the MD equilibration step. The poses are numbered from the best-scoring (Pose 1) to worst-scoring (Pose 5) according to Vina. The last data row gives the computed binding free energies (kcal/mol) starting from each pose or the crystal structure. Uncertainties are provided in parentheses, according to Eq. 15. The tabulated double decoupling free energies were computed using TI with 12-point Gaussian quadrature; the other free energy terms were computed as shown in Table 1.

|  | Starting Structure |  |  |  |  |  |
| --- | --- | --- | --- | --- | --- | --- |
|  | Crystal | Pose 1 | Pose 2 | Pose 3 | Pose 4 | Pose 5 |
|  | RMSD from crystal structure (Å) |  |  |  |  |  |
| Initial | 0.00 | 5.36 | 2.00 | 5.58 | 5.50 | 1.07 |
| Equilibrated | 1.30 | 5.26 | 0.45 | 5.33 | 4.23 | 0.74 |
|  | Computed binding free energy (kcal/mol) |  |  |  |  |  |
| $-\Delta G_{bind}^o$ | 6.10 (0.58) | 2.53 (0.85) | 6.72 (0.62) | 2.61 (0.75) | 1.55 (0.58) | 6.53 (0.77) |

The free energies reported in Table 2 used the TI-GQ method to compute the decoupling and annihilation terms, but we also ran these calculations with MBAR, and the two methods should give the same result in the limit of infinite sampling and infinitesimally spaced windows. Table 3 compares the results from these methods for each of the four decoupling or annihilation terms (Equations 13 and 14), for the crystal structure calculations represented in Table 2. The TI-GQ calculations used 12 windows, while MBAR used 23 windows between *λ* = 0 and *λ* = 1 (see SI). The number of steps for each window was the same in both methods, resulting in nearly twice as much simulation time for the MBAR calculations. The deviations between the two sets of calculations range from 0.09 to 0.36 kcal/mol for the four terms, showing good consistency between the two approaches. Note that the TI-GQ and MBAR calculations use independent sets of data, since the lambda values, and thus the simulation input parameters for each window, are not the same between the two. Table 3 also shows lower reported uncertainties for MBAR, when compared to TI-GQ. This could be due to the MBAR calculations having a greater number of lambda values between 0 and 1, and thus more sampling between the coupled and decoupled states.

**Table 3:** Comparison of annihilation/decoupling free energies (kcal/mol) computed with MBAR and TI-GQ. Estimated uncertainties are given in parenthesis.

|  | MBAR | TI-GQ | Difference |
| --- | --- | --- | --- |
| $\Delta G_{elec,bound}$ | -8.67 (0.26) | -8.31 (0.45) | -0.36 |
| $\Delta G_{LJ,bound}$ | 10.14 (0.20) | 9.99 (0.22) | 0.15 |
| $-\Delta G_{LJ,unbound}$ | 0.94 (0.08) | 1.00 (0.11) | -0.06 |
| $-\Delta G_{elec,unbound}$ | 11.09 (0.02) | 11.00 (0.12) | 0.09 |
| $\Delta G_{trans}$ | 13.50 (0.34) | 13.68 (0.53) | -0.18 |

The present results illustrate the potential for physics-based, absolute binding free energy methods to distinguish accurate from inaccurate poses and to compute standard binding free energies that may be compared directly with experiment. Absolute binding free energy methods are particularly suitable for virtual screening, where the compounds of interest can be highly diverse. In contrast, relative binding free energy methods,^8–12^ which involve al-chemically changing one compound to another, are best suited to for comparing the affinities of chemically similar compounds, as in the scenario of lead optimization. In summary, although more testing is clearly needed, this initial study is encouraging and motivates future broader tests on other protein-ligand systems.

### 5.2 Performance and costs

All simulations in this work used AMBER18’s *pmemd*.*cuda* software, which provides remarkable performance on GPUs.^30,61^ Detailed timings for the various components of the DD calculations on a single NVIDIA GTX 1070 GPU, for a single ligand pose, are provided in Table 4. Each calculation involved a total of 1.24 *µ*s of simulations, which took about seven days on an otherwise unoccupied machine. This wall-clock time can be easily reduced by trivial parallelization across GPUs, because all of the free energy windows can be simulated independently. With full parallelization along these lines, the minimal wall-clock time becomes the computational time needed for the longest simulations in the free energy calculations, which are close to six hours for the ligand LJ decoupling windows in the binding site. We expect these times to be nearly halved on a more recent model NVIDIA RTX 2080 GPU.^61^

**Table 4:** Computational speed and timings (wall clock) to run a binding free energy calculation with a single GTX 1070 GPU on a computer with no other calculations running.

| Stage | Atoms | Speed | Simulation time (ST) | Computation time (CT) | ST per window | CT per window |
| --- | --- | --- | --- | --- | --- | --- |
| Free energy terms |  |  |  |  |  |  |
| Equilibration/<br>Preparation | $\sim 40k$ | $\sim 205$ ns/d | 80ns | 0.39 d | - | - |
| $\Delta G_{p,att}$ | $\sim 40k$ | $\sim 205$ ns/d | 96ns | 0.47 d | 6ns | 0.70 h |
| $\Delta G_{l,conf,att}$ | $\sim 40k$ | $\sim 205$ ns/d | 96ns | 0.47 d | 6ns | 0.70 h |
| $\Delta G_{l,TR,att}$ | $\sim 40k$ | $\sim 205$ ns/d | 96ns | 0.47 d | 6ns | 0.70 h |
| $\Delta G_{elec,bound}$ | $\sim 40k$ | $\sim 95$ ns/d | 28.8ns | 0.30 d | 2.4ns | 0.61 h |
| $\Delta G_{LJ,bound}$ | $\sim 40k$ | $\sim 95$ ns/d | 288ns | 3.03 d | 24ns | 6.10 h |
| $\Delta G_{LJ,unbound}$ | $\sim 6k$ | $\sim 490$ ns/d | 144ns | 0.29 d | 12ns | 0.59 h |
| $\Delta G_{elec,unbound}$ | $\sim 6k$ | $\sim 450$ ns/d | 28.8ns | 0.06 d | 2.4ns | 0.13 h |
| $\Delta G_{l,conf,rel}$ | $\sim 6k$ | $\sim 860$ ns/d | 192ns | 0.22 d | 12ns | 0.33 h |
| $\Delta G_{p,rel}$ | $\sim 40k$ | $\sim 205$ ns/d | 192ns | 0.94 d | 12ns | 1.40 h |
| Total time |  |  |  |  |  |  |
| $\Delta G_{bind}^{\circ}$ | - | - | 1.24 $\mu s$ | 6.64 d | - | - |

It is also worth noting that the speed of a simulation with *pmemd*.*cuda* depends mainly on the choice of GPU, and far less on the choice of CPU, motherboard, etc. As a consequence, high-performance simulations can be achieved at low cost by using computers configured with strong GPUs but minimalist components otherwise. As an example, the present SI suggests a machine build using affordable components, with 2 GTX 1070 GPUs, capable of running a calculation like the ones presented here in a few days. The final cost, including peripherals, is ∼$1520, or only $760 per installed GPU, and could be further reduced by choosing more modest components. We also present the option of using the newer RTX 2080 instead of the GTX 1070, showing that the price/performance ratio can be preserved or even improved by that choice.

## 6 Ready for high-throughput?

Our main goal in creating the BAT.py software is to enable rigorous ABFE calculations in a high-throughput regime and thus to enable their use in the first stages of drug discovery. To make that a reality, we believe two things are indispensable: automation of the free energy calculations and high performance of the simulations.

The first requirement is made possible by the workflow of BAT.py (Fig. 1), which can start either from a set of docked poses of a given ligand or from a crystal structure. Once the necessary input parameters are determined and optimized for a new receptor, along with a docking procedure to generate plausible poses, the calculations can be performed for a library of ligands by simply running BAT.py in the command line for each compound. The flexibility of BAT.py allows one to choose the optimal force-field parameters, adjust the simulation times to prioritize speed or exhaustive sampling, and set the most suitable protein conformational restraints. The second requirement is that the simulations can be performed with high performance and at low cost. This capability is now in view, given the fast *pmemd*.*cuda* implementation of AMBER and GPU implementations of other simulation packages, coupled with the increasing performance and affordability of GPUs.

Thus, these two conditions are now largely satisfied. This makes it possible to move to expanded testing of ABFE calculations as a tool for scoring docked ligand poses, while also estimating overall binding free energies, so that many ligands can be ranked in terms of their calculated affinities. We plan next to extend these calculations to other protein systems, testing different force fields, water models and simulation parameters, including in the context of the rolling CELPP pose-prediction exercise. ^55^

## Supporting information

BAT.py User Guide

## Acknowledgement

GH thanks FAPESC and CNPq for the research grants. MKG acknowledges funding from National Institute of General Medical Sciences (Grant number GM061300). The contents of this paper are solely the responsibility of the authors and do not necessarily represent the official views of the NIH. MKG has an equity interest in and is a cofounder and scientific advisor of VeraChem LLC.

