## Supporting Info for "Automated docking refinement and virtual compound screening with absolute binding free energy calculations"

### 1 Input files

#### 1.1 Docked poses with TI-GQ

Below are the input file options used to process the five poses obtained from the docking procedure, as explained in the main text, using the TI-GQ method for the double decoupling calculations. The distances are in Å, the force constants for distances are in kcal/(mol.Å$^2$), and the force constants for angles and dihedrals are in kcal/(mol.rad$^2$).

```
calc_type = dock

celpp_receptor = LMCSS-5uf0_5uez

poses_list = [0,1,2,3,4]

ligand_name = LIG

P1 = :403@CA

P2 = :424@CA

P3 = :450@CA

fe_type = dd-rest

components = [ e v w f t r l a c ]

release_eq = [ 5.00 2.50 1.00 0.50 0.20 0.10 0.05 0.02 0.00 ]

attach_rest = [ 0.00 0.20 0.40 0.80 1.60 2.40 4.00 5.50 8.65 11.80
   18.10 24.40 37.00 49.60 74.80 100.00 ]

lambdas = [ 0.00922 0.04794 0.11505 0.20634 0.31608 0.43738 0.56262
   0.68392 0.79366 0.88495 0.95206 0.99078 ]

pull_ligand = no

weights = [ 0.02359 0.05347 0.08004 0.10158 0.11675 0.12457 0.12457
   0.11675 0.10158 0.08004 0.05347 0.02359 ]

dd_type = TI

blocks = 5

rec_distance_force = 10.0

rec_angle_force = 250.0
```

```
rec_dihcf_force = 50.0
rec_discf_force =  5.0
lig_distance_force = 5.0
lig_angle_force = 250.0
lig_dihcf_force = 70.0
lig_discf_force =  5.0
water_model = TIP3P
num_waters  = 13000
buffer_x    = 12
buffer_y    = 12
lig_buffer  = 15
neutralize_only = no
cation = Na+
anion = Cl-
num_cations = 32
num_cat_ligbox = 8
hmr         = yes
Temperature = 298.15
eq_steps1   = 500000
eq_steps2   = 15000000
prep_steps1 = 1000000
a_steps1    = 500000
a_steps2    = 1000000
l_steps1    = 500000
l_steps2    = 1000000
t_steps1    = 500000
t_steps2    = 1000000
c_steps1    = 1000000
c_steps2    = 2000000
```

```
r_steps1     = 1000000
r_steps2     = 2000000
e_steps1     = 200000
e_steps2     = 400000
v_steps1     = 2000000
v_steps2     = 4000000
w_steps1     = 1000000
w_steps2     = 2000000
f_steps1     = 200000
f_steps2     = 400000
rec_bb       = yes
bb_start     = 379
bb_end       = 389
bb_equil     = yes
l1_x         = 0.00
l1_y         = -6.25
l1_z         = 8.50
l1_zm        = 20.0
l1_range     = 2.50
min_adis     = 4.00
max_adis     = 8.00
ntpr = 1000
ntwr = 1000
ntwe = 0
ntwx = 2500
cut = 9.0
gamma_ln = 1.0
barostat = 2
dt = 0.004
```

```
receptor_ff = protein.ff14SB

ligand_ff = gaff
```

#### 1.2 Crystal structure with TI-GQ

For the 5uf0 crystal structure calculations, the following parameters are changed relative to those in Section 1.1:

```
calc_type = crystal

celpp_receptor = 5uf0

ligand_name = 89J
```

#### 1.3 Double decoupling with MBAR

When applying the MBAR[S1] procedure for the double decoupling calculations, instead of TI-GQ, the following parameters are changed relative to Section 1.1:

```
lambdas = [ 0.0001 0.02 0.04 0.06 0.08 0.10 0.15 0.20 0.25 0.30 0.40
   0.50 0.60 0.70 0.75 0.80 0.85 0.90 0.92 0.94 0.96 0.98 0.9999 ]

dd_type = MBAR
```

### 2 Free energy components

Table S1: All free energy terms and final binding free energies for each of the docked poses and the crystal structure, generated with the TI-GQ method with double decoupling. The two top rows give the ligand RMSD in the binding site relative to the 5uf0 structure, for the docked poses and crystal structure systems before (Initial) and after equilibration (Equilibrated). Uncertainties ($\sigma$) are shown in parentheses, with all energies in kcal/mol.

| System | Crystal | pose 1 | pose 2 | pose 3 | pose 4 | pose 5 |
|---|---|---|---|---|---|---|
| RMSD from crystal structure (Å) | | | | | | |
| Initial | 0.00 | 5.36 | 2.00 | 5.58 | 5.50 | 1.07 |
| Equilibrated | 1.30 | 5.26 | 0.45 | 5.33 | 4.23 | 0.74 |
| Free energy terms | | | | | | |
| $\Delta G_{p,att}$ | 15.23 (0.11) | 15.85 (0.34) | 15.71 (0.23) | 15.66 (0.27) | 16.47 (0.26) | 15.97 (0.12) |
| $\Delta G_{l,conf,att}$ | 8.89 (0.06) | 9.96 (0.16) | 8.67 (0.15) | 8.81 (0.09) | 11.48 (0.17) | 9.00 (0.04) |
| $\Delta G_{l,TR,att}$ | 7.30 (0.05) | 5.18 (0.21) | 5.27 (0.04) | 7.17 (0.12) | 7.98 (0.26) | 5.32 (0.13) |
| $\Delta G_{elec,bound}$ | -8.31 (0.45) | -11.64 (0.23) | -7.65 (0.26) | -12.96 (0.11) | -12.99 (0.20) | -9.34 (0.35) |
| $\Delta G_{LJ,bound}$ | 9.99 (0.22) | 11.00 (0.68) | 11.33 (0.45) | 10.19 (0.64) | 6.83 (0.33) | 11.94 (0.64) |
| $-\Delta G_{LJ,unbound}$ | 1.00 (0.11) | 0.93 (0.07) | 1.00 (0.10) | 0.97 (0.15) | 0.88 (0.12) | 1.09 (0.09) |
| $-\Delta G_{elec,unbound}$ | 11.00 (0.12) | 11.15 (0.06) | 11.25 (0.09) | 11.23 (0.13) | 11.48 (0.07) | 11.21 (0.12) |
| $\Delta G_{l,TR,rel}$ | -12.77 | -12.64 | -12.96 | -12.65 | -12.64 | -12.88 |
| $\Delta G_{l,conf,rel}$ | -8.99 (0.07) | -10.74 (0.07) | -9.39 (0.08) | -9.30 (0.10) | -11.44 (0.06) | -9.27 (0.05) |
| $\Delta G_{p,rel}$ | -17.23 (0.19) | -16.51 (0.06) | -16.51 (0.06) | -16.51 (0.06) | -16.51 (0.06) | -16.51 (0.06) |
| Final binding free energies | | | | | | |
| $-\Delta G^{\circ}_{bind}$ | **6.10 (0.58)** | **2.53 (0.85)** | **6.72 (0.62)** | **2.61 (0.75)** | **1.55 (0.58)** | **6.53 (0.77)** |

### 3 Using the Attach-Pull-Release Method

#### 3.1 Input variables

In order to use APR, some options have to be either changed or added, relative to a BAT input file for a DD calculation (Section 1).

```
fe_type = pmf

translate_apr = [ 0.00 0.40 0.80 1.20 1.60 2.00 2.40 2.80 3.20 3.60
    4.00 4.40 4.80 5.20 5.60 6.00 6.40 6.80 7.20 7.60 8.00 8.40 8.80
    9.20 9.60 10.00 10.40 10.80 11.20 11.60 12.00 12.40 12.80 13.20
    13.60 14.00 14.40 14.80 15.20 15.60 16.00 ]
```

```
pull_ligand = yes
u_steps1    = 5000000
u_steps2    = 20000000
prep_steps2 = 100000
pull_spacing = 0.1
```

#### 3.2 Theory and Method

When using APR, the transfer free energy $\Delta G_{trans}$ (Fig. 2 and Eq. 3 from manuscript) is computed as the reversible work of pulling the restrained ligand from the restrained binding site and into bulk solvent, as previously described.[S2]

$$\Delta G_{trans} = \Delta G_{pull} \tag{1}$$

This is done through a series of windows between the bound state and the state with the ligand in bulk solvent. The reaction coordinate of the APR process is D1, the distance between the first dummy atom N1 and the first ligand anchor atom L1. The physical path defined by this distance is broken into windows, each with a harmonic umbrella restraint having a different value of the reference distance, $r_0$ (Eq. 11 from manuscript). The location of each of the pulling windows, as well as the pulling force constant, can be defined in the BAT.py input file using the *translate_apr* and *lig_distance_force* variables, respectively.

The restraint free energy terms when using APR and DD for the same system are the identical, except for the release of the ligand TR restraints in bulk to the standard concentration $C^{\circ}$, $\Delta G_{l,TR,rel}$ (Eq. 10 from manuscript). APR has a different value for $r_0$ in this equation, namely the final pulling distance of the umbrella sampling process, which is the last value in the *translate_apr* array. In the current example, the umbrella windows go from from 5.0 Å to 21 Å in 0.4 Å increments.

##### 3.3 Preparation

The initial state for each of the umbrella windows is obtained by a process similar to steered molecular dynamics (SMD), which is performed during the preparation stage when the *pull_ligand* variable is activated. This causes the restrained ligand to be brought from the binding site to bulk solvent through a sequence of simulations with all restraints activated, each adding a small increment (e.g. 0.1 Å) to the reference value, $r_0$, of the D1 distance restraint. The increment size is defined by the *pull_spacing* variable, and the initial and final SMD distances are defined by the first and last elements of the *translate_apr* array. The number of steps for each simulation is specified with *prep_steps2*. The final states drawn from a subset of this SMD process (e.g. every 0.4 Å) will be used as the initial states of the umbrella windows along the pulling pathway.

##### 3.4 Free energy calculation

By analogy with Table 1 in the main text, the letter associated with the umbrella sampling component along the pulling pathway is **u**, with *u_steps1* and *u_steps2* the variables defining the number of equilibrium and production steps of each window. The MBAR method is used to obtain the reversible work of pulling the ligand from the binding site to bulk solvent, which corresponds to the $\Delta G_{pull}$ term.

In Table S2 we show the results of the automated APR calculation for the 5uf0 crystal structure. All restraint free energies are the same as the ones from the DD calculations for this system (Table S1), except for the analytical release $\Delta G_{l,TR,rel}$, as explained above. The binding free energy calculated using APR, -5.58 kcal/mol, is within numerical uncertainty when compared to the DD result for this system, which is -6.10 kcal/mol. This APR calculation took a total of 4.1 $\mu$s of simulation time, which may be compared with 1.2 $\mu$s for the DD calculations detailed in the main text.

Table S2: All free energy terms and final binding free energy for an APR calculation of the binding free energy of the 5uf0 crystal structure. Uncertainties are shown in parentheses, with all energies in kcal/mol.

| Free energy terms | |
|---|---|
| $\Delta G_{p,att}$ | 15.23 (0.11) |
| $\Delta G_{l,conf,att}$ | 8.89 (0.06) |
| $\Delta G_{l,TR,att}$ | 7.30 (0.05) |
| $\Delta G_{pull}$ | 11.45 (0.62) |
| $\Delta G_{l,TR,rel}$ | -11.07 |
| $\Delta G_{l,conf,rel}$ | -8.99 (0.07) |
| $\Delta G_{p,rel}$ | -17.23 (0.19) |
| Final binding free energy | |
| $\boldsymbol{-\Delta G^{\circ}_{bind}}$ | **5.58 (0.67)** |

### 4 Sample hardware setup

Table S3 details a complete machine built with two NVIDIA GTX 1070 GPUs and relatively cheap other components. This setup is able to run independent simulations on each GPU with the same performance as shown in Table 4 of the main text. A similar setup can be built replacing the GTX 1070 graphics card with newer cards having the Turing architecture, such as the NVIDIA RTX 2080, also shown in Table S3. Even though this is more costly, it will roughly double the speed of the calculations and thus preserve or improve the price/performance ratio.[S3]

Table S3: Prices of parts and full machine built to run simulations using either NVIDIA GTX 1070 or RTX 2080 graphics processing units. Prices of a large online retailer as of April/2020, given in US dollars.

| Hardware | Quantity | Total Price |
|---|---|---|
| Motherboard ASUS TUF Z370 LGA1151 DDR4 | 1 | $150 |
| Intel Core i5-8400 6 Cores 4.0GHz LGA1151 | 1 | $190 |
| Corsair RM850x 850W 80+ Gold Power Supply | 1 | $150 |
| EVGA GeForce GTX 1070 SC GAMING | 2 | $800 |
| Gigabyte GeForce RTX 2080 OC GAMING | 2 | $1400 |
| Samsung 860 EVO 500GB 2.5 Inch SATA III SSD | 1 | $90 |
| Corsair Vengeance LPX 16GB DDR4 3000MHz | 1 | $90 |
| Rosewill TYRFING ATX Computer Case | 1 | $50 |
| Full workstation | | |
| With 2 GTX 1070 GPUs | | $1520 |
| With 2 RTX 2080 GPUs | | $2120 |
| Price per GTX 1070 including peripherals | | $760 |
| Price per RTX 2080 including peripherals | | $1060 |
