## Supplementary material for "Automated docking refinement and virtual compound screening with absolute binding free energy calculations": BAT.py User Guide

### User guide for BAT.py – v1.0

#### 1. Introduction

The Binding Affinity Tool (BAT.py) is designed to fully automate absolute binding free energy calculations, starting only from a crystal structure or a docked complex. The building of the simulation boxes, generation of all the needed parameters, set up of the various simulation windows, running the simulations, and the final free energy analysis are all done without any manual interference. BAT.py uses the *pmemd.cuda* sofware from AMBER [1], which has shown very high performance on Graphics Processing Units (GPUs) at a reduced cost (https://ambermd.org/GPUPerformance.php). We believe that our implementation can be used for high-throughput search of high-affinity ligands to a given receptor, using a rigorous physics-based free energy approach. BAT.py can also be applied for parameter testing and optimization, searching for the optimal ones to be used on a given system. The program, along with a tutorial and input files for various protein systems, can be found on the link https://github.com/Gheinzelmann/BAT.py.

In this user guide we will first describe the workflow of the program, then the various components of the free energy calculation, and how the simulations are analyzed in order to obtain the quantities of interest. All the parameters needed for the program input file, and how they apply to the various calculation steps, will also be described in detail. Finally, we will explain how to add a new protein system to our automated protocol.

#### 2. Workflow

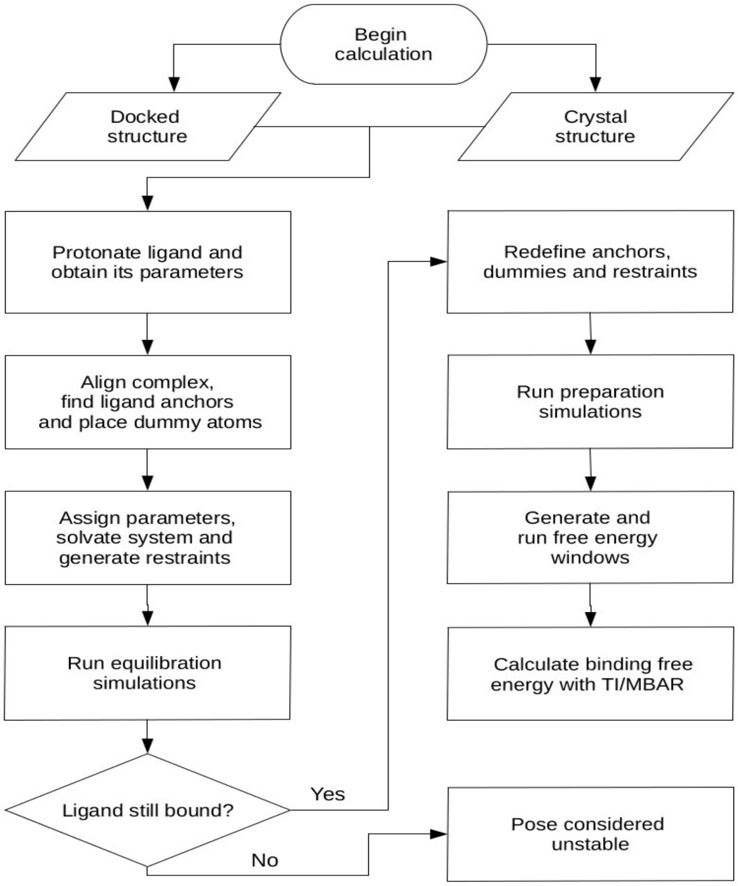

The workflow above shows the equilibration, preparation and calculation procedures. It begins by first building the initial complex, either starting from the docked receptor and ligand files or directly from the receptor-ligand co-crystal structure. The necessary ligand parameters are obtained using Antechamber [2], with the General Amber Force-Field (GAFF or GAFF2) for the bonded and LJ parameters [3], and the AM1-BCC model for the partial atomic charges [4,5]. The system is then aligned to a reference structure of a similar protein using the program MUSTANG [6], so that the ligand anchors and dummy atom coordinates can be automatically assigned.

With all the coordinates of the initial system already set, the complex is then placed in a water box with a given ion concentration, and the necessary restraints are applied. An initial equilibration is then performed, with a restrained receptor, and the translational/rotational restraints of the ligand being gradually released in order to find a nearby energy minimum. At the end of this last step, the ligand might still be bound or it might have left the binding site in the case of unstable binding mode. If the latter happens, this pose is considered unstable and no further simulations are performed for this system.

If the ligand is still bound after equilibration, then the preparation of the system for the binding free energy calculations is performed. The preparation starts from the last state from equilibrium, reassigning the ligand anchors, repositioning the dummy atoms, solvating/ionizing the system and and redefining the restraints. This is necessary since the unrestrained ligand can adopt a different binding mode in the last stage of the equilibration step, which requires a new reference state for the free energy calculations. The preparation simulations may involve the pulling of the ligand from the binding site to bulk, if APR is to be used, or only the simulation of the restrained ligand in the bound state, if only DD is employed.

Starting from the last state of the preparation step, all the necessary simulation windows are now created for the binding free energy calculation. They involve various components for the application/removal of restraints, as well as the pulling/decoupling of the ligand, which are explained in detail in the next section. Once the free energy simulations are concluded, they can be analyzed using the Multistate Bennett Acceptance Ratio (MBAR) [7], Thermodynamic Integration with Gaussian Quadrature (TI), or analytically, depending on the stage and the choice in the input file.

#### 3. Free Energy Components

The BAT.py expression for the calculated binding free energy is defined as follows:

$$-\Delta G_{bind}^{o} = \Delta G_{p,att} + \Delta G_{l,conf,att} + \Delta G_{l,TR,att} + \Delta G_{trans} + \Delta G_{l,TR,rel} + \Delta G_{l,conf,rel} + \Delta G_{p,rel}$$

In the equation above, the *att* index denotes attachment of restraints in the bound state, and *rel* indicates release of restraints with the ligand in bulk. The *l* and *p* indexes are for ligand and protein (receptor), respectively, *conf* is for conformational restraints and *TR* is for translational/rotational restraints. The $\Delta G_{trans}$ term is the free energy of transferring of the ligand from the receptor binding site to bulk with all restraints applied, using either a physical reaction coordinate (APR), or an alchemical transformation (DD):

$$\Delta G_{trans-APR} = \Delta G_{pull} \qquad \textbf{(APR)}$$

$$\Delta G_{trans-DD} = \Delta G_{dcpl,elec,bound} + \Delta G_{dcpl,LJ,bound} - \Delta G_{dcpl,elec,unbound} - \Delta G_{dcpl,LJ,unbound} \qquad \textbf{(DD)}$$

In the case of APR, $\Delta G_{trans\text{-}APR}$ is equal to the pulling free energy of the ligand from the binding site to bulk, which is done using umbrella sampling, as in Ref [8]. For the double decoupling procedure, $\Delta G_{trans\text{-}DD}$ is equal to the sum of four terms, as shown in the equation above. The index *dcpl* stands for decoupling, *elec* for electrostatic interactions and *LJ* for Lennard-Jones interactions, with these calculations being performed with the ligand both in the binding site (*bound)* and in bulk (*unbound*).

Table I summarizes all the free energy components from our simulations, with each identified by a letter:

| Description | Letter | System | Free Energy Method | Free energy term |
|---|---|---|---|---|
| Attachment of receptor conformational restraints | **a** | Complex | MBAR | $\Delta G_{p,att}$ |
| Attachment of ligand conformational restraints | **l** | Complex | MBAR | $\Delta G_{l,conf,att}$ |
| Attachment of ligand TR restraints | **t** | Complex | MBAR | $\Delta G_{l,TR,att}$ |
| Pulling stage (umbrella sampling) | **u** | Complex* | MBAR | $\Delta G_{pull}$ |
| Decoupling of ligand charge interactions (binding site) | **e** | Complex | MBAR/TI | $\Delta G_{dcpl,elect,bound}$ |
| Decoupling of ligand LJ interactions (binding site) | **v** | Complex | MBAR/TI | $\Delta G_{dcpl,LJ,bound}$ |
| Decoupling of ligand charge interactions (bulk) | **f** | Ligand only | MBAR/TI | $\Delta G_{dcpl,elec,unbound}$ |
| Decoupling of ligand LJ interactions (bulk) | **w** | Ligand only | MBAR/TI | $\Delta G_{dcpl,LJ,unbound}$ |
| Release of ligand TR restraints | **b** | Ligand only | Analytical | $\Delta G_{l,TR,rel}$ |
| Release of ligand conformational restraints | **c** | Ligand only | MBAR | $\Delta G_{l,conf,rel}$ |
| Release of receptor conformational restraints | **r** | Receptor only | MBAR | $\Delta G_{p,rel}$ |

* The receptor and ligand will be physically separated during the pulling simulations

When the calculations are set up, the windows from each free energy component will have their corresponding letter followed by the window number, starting at 0. The number of windows and their properties can be defined in the input file. The letters also identify the free energy output files, which are stored in the ./data folder of each component, after the analysis is performed. More information on the nature of each of the restraints, and the free energy methods we use, can be found in Refs. [8,9].

#### 4. Input file

Various options concerning the creation of the systems, simulations and analysis, can be chosen in the input file:

`calc_type:` Accepts the options "dock" or "crystal", for a receptor ligand pair of pdb files, or a complex co-crystal structure, respectively. The system initial pdb files should be located in the ./all-poses folder.

`celpp_receptor`: Sets the name of the receptor, followed by _docked. For example, choose “LMCSS-5uf0_5uez” for a receptor file called LMCSS-5uf0_5uez_docked.pdb. The naming is based on the CELPP challenge procedure. For a crystal structure, put the four letter identifier for the structure, for example “5uf0” for the 5uf0.pdb crystal structure.

`poses_list`: The list of poses that will be used for the calculations. The list should be placed in brackets ans separated by commas. Ex: “[0,1,2,3,4]”. The docked poses files in the ./all-poses folder must be listed accordingly as pose0.pdb, pose1.pdb, pose2.pdb, etc. This parameter is not used for crystal structure calculations.

`ligand_name`: The ligand residue name in the docked poses above, if `calc_type` is set to “dock” (Ex: “LIG”), or in the initial crystal structure, if `calc_type` is set to “crystal” (Ex: “89J” for the 5uf0 ligand.).

`P1, P2` and `P3`: These define the anchor atoms of the receptor, which have to be determined beforehand. The original protein sequence numbering should be used here, using AMBER masks to define each atom. Ex: “:403@CA” for the CA atom of residue 403.

`fe_type`: Designates which components are to be included in the free energy calculations. For example, if all the APR and DD calculations are to be performed, choose “all” for this option. If only double decoupling with restraints will be performed, choose “dd-rest”, and if only APR choose “pmf-rest”. The “custom” option allows to choose any combination of components, using the `components` option below.

`components`: If the option “custom” is set in the option above, choose the components you want to calculate, using a list of letters separated by spaces inside a bracket. Ex: “[ c l t a ]”.

`release_eq`: The weights for the gradual release of the restraints in the equilibrium stage, going from 100 (fully restrained) to 0 (unrestrained). Each option will be a new simulation, and they are performed in sequence. Use a list of letters separated by spaces inside a bracket to define these weights. Ex: “[ 5.00 2.50 1.00 0.00 ]”. A single 0.00 inside the brackets (Ex: “[ 0.00 ]”) will run just one equilibrium simulation without any ligand restraints.

`attach_rest`: List of weights for the spring constant of each window during the attaching/releasing of restraints using MBAR (components **a**, **l**, **t**, **c** and **r**). The total number of windows for each of these components will be the size of the array. Ex: “[ 0.00 2.00 4.00 16.00 64.00 100.00 ]” for a total of 6 windows.

`translate_apr`: Windows for the umbrella sampling (pulling) procedure of APR, identified by the letter **u**. It starts from 0.00 (bound state) until the desired reference distance between the receptor and the ligand in the unbound state. The number of windows is the size of the array. Ex: “[ 0.00 1.00 2.00 3.00 4.00 5.00 ]” for a total of 6 windows, ending 5.00 Å away from the initial reference distance between N1 and L1, which is 5.00 Å by construction.

`lambdas`: Lambda values for the double decoupling procedure, going from 0.00 to 1.00. Ex: For a 12-point Gaussian quadrature, choose “[ 0.00922 0.04794 0.11505 0.20634 0.31608 0.43738 0.56262 0.68392 0.79366 0.88495 0.95206 0.99078 ]” for the lambda array values.

`pull_ligand`: Choice to pull the ligand from the binding site or not during preparation. This is needed for the APR method, but not needed for double decoupling. Choose “yes” or “no” for this option.

`pull_spacing`: The interval between each ligand position during the preparation simulations, if the option above is set to yes. The final distance is the last value in the translate_apr array. Ex: “0.1” for a pulling interval of 0.1 Å.

rec_distance_force: Distance spring constant for the receptor rigid body restraints, identified as D2 in Ref. [8]. Use units of kcal/mol.Å$^2$.

rec_angle_force: Angle and torsion angle spring constants for the receptor rigid body restraints, identified as A3, A4, T4, T5 and T6 in Ref. [8]. Use units of kcal/mol.rad$^2$. The forces from rec_distance_force and rec_angle_force are included to keep the receptor in the laboratory reference frame.

rec_dihcf_force: Final spring constant for the protein conformational dihedral restraints, if this option is activated (see rec_bb variable). The nature of these restraints, and how they are implemented, are explained in Ref. [9]. Use units of kcal/mol.rad$^2$.

rec_discf_force: Final spring constant for the protein conformational distance restraints between the anchor atoms. Use units of kcal/mol.Å$^2$.

lig_distance_force, lig_angle_force, lig_dihcf_force and lig_discf_force: Final spring constants for the ligand restraints, defined the same way as the receptor above. The value of lig_distance_force also designates the spring constant used during the pulling simulations. The nature of the ligand conformational restraints, and how they are implemented, are explained in Ref. [9].

water_model: The water model used in the calculations. Supported options are “TIP3P”, “TIP4PEW” and “SPCE”.

num_waters: Number of waters used in the simulations of the complex and the *apo* protein.

buffer_x and buffer_y: Options for the water padding in the *x* and *y* axes of the system, remembering that the pulling is done along the *z* coordinate. The dependent variable is the padding in the *z*-axis, so make sure you have enough waters to cover the complex, and also allow the pulling of the ligand if APR is used.

lig_buffer: Water padding in the three Cartesian axes for the box with only the ligand in it.

neutralize_only: Option to add ions only to neutralize the system, or to also include an additional number of ions. Accepts options “yes” or “no”.

cation and anion: Cation and anion species to be used, accepts all ions supported by the Joung and Cheatham monovalent ion parameters [10]. Ex: “Na+” and “Cl-”.

num_cations: Number of cations to be added after neutralization, for the desired ion concentration, for simulations of the complex and the *apo* protein. The number of anions is the dependent variable, since the systems are always neutral.

num_cations_ligbox: Number of cations to be added after neutralization, for the desired ion concentration, for the smaller ligand box.

hmr: Use hydrogen mass repartitioning [11] or not. Accepts options “yes” and “no”.

temperature: Temperature of the simulated systems, in Kelvin (K).

eq_steps1: Number of steps for each simulation of the gradual release of restraints, during the equilibration procedure.

eq_steps2: Number of steps for the last simulation of the equilibration procedure, in which the ligand is unrestrained.

prep_steps1: Number of steps for the first simulation of the preparation step, in which the ligand is fully restrained in the bound state.

prep_steps2: Number of steps for each of the pulling simulations, during the preparation procedure, if pull_ligand is set to yes. The distance separation of each of these steps is defined in the pull_spacing option.

[component]_steps1: Number of steps of equilibration, for each window of the various components of the free energy calculation, with the component letters shown in Table I. No data is collected during this simulation.

[component]_steps2: Number of steps for the production stage of each window of the various components of the free energy calculation, in which data is collected.

rec_bb: Choice to use or not protein (receptor) backbone dihedral restraints, accepting "yes" or "no".

bb_start and bb_end: If the option above is activated, these variables define the residue range of the protein backbone dihedral restraints, using the original protein sequence numbering.

bb_equil: Choice to keep the protein backbone restraints fully attached during the ligand equilibration, or leave it unrestrained so it can adapt to the docked ligand. If the latter is chosen, the reference coordinates for the backbone restraints come from the final state of equilibration. Accepts "yes" or "no".

l1_x: distance in the *x* axis between the first protein anchor atom P1 and the center of the "strike zone" for the search of the ligand first anchor L1. More details on this procedure can be found in section 6 of this guide and also in Ref. [9].

l1_y: Same as the previous one, but for the *y* axis distance.

l1_z: Minimum distance between the first protein anchor atom P1 and the first ligand anchor L1, in the *z* axis.

l1_zm: Maximum distance between the first protein anchor atom P1 and the first ligand anchor L1, in the *z* axis.

l1_range: Size of the "strike zone" for the first ligand anchor atom search, which is a square with sides having twice the value of this parameter (2*l1_range).

min_adis and max_adis: Minimum and maximum distance between the ligand anchors.

dd_type: Type of integration method for the decoupling components of the binding free energy calculation (e, v, f and w). If "TI" is chosen, Gaussian quadrature is applied, if "MBAR" is chosen, the latter is used to calculate these components. Remember that the lambda values have to be suitable for either type of integration method.

`weights`: Weights for Gaussian quadrature calculations, in case the TI option is chosen above. These values must correspond to the values in the `lambdas` array, for the procedure to be applied correctly. In the case of a 12-point Gaussian quadrature, write "[ 0.02359 0.05347 0.08004 0.10158 0.11675 0.12457 0.12457 0.11675 0.10158 0.08004 0.05347 0.02359 ]" for this variable.

`blocks`: Number of blocks for block data analysis. This separates the simulation data in blocks and provides the results for each, so the temporal variation and convergence of the results can be assessed. This option is also used for the calculation of the uncertainties of each free energy component [9].

`ntpr`, `ntwr`, `ntwe`, `ntwx`, `cut`, `gamma_ln`, `barostat` and `dt`: Options for running the various simulations, such as output frequency, non-bonded cutoff, barostat type, time step, and others. These use the same variables as the ones from the *pmemd.cuda* simulation input file, and their definitions can be found in the AMBER user guide.

`receptor_ff`: Choice of force field for the receptor atoms, such as "protein.ff14SB" or "protein.fb15". Also supports parameters for DNA and RNA, such as "DNA.OL15" and "RNA.OL3".

`ligand_ff`: Choice of force field for the ligand Lennard-Jones and bonded parameters. Accepts either "gaff" or "gaff2". The ligand partial charges are always determined by the AM1-BCC model.

#### 5. Adding new ligands to a given protein

In the example provided in the BAT folder, binding free energy calculations are performed for the 5uf0 crystal structure, as well as 5 docked poses of the same ligand docked to the receptor with the 5uez structure. The protein receptor is the second bromodomain of the BRD4 protein – BRD4(2). The ./all-poses folder in this example contains the original 5uf0.pdb file, as well as a set of pdb files for the docked poses and receptor. The same procedure can be applied to any ligand that binds to BRD4(2), as explained below:

**5.1 Docked complexes:** In order to generate a set of pdb files for the docked poses and receptor, there are two options, either do a manual docking with chosen input files, or using the CELPP challenge workflow. Both options use AutoDockTools [12], Chimera [13], and AutoDock Vina [14] to prepare the files and run the docking, so you must have them in your path in order to perform this stage. For the first option, the ./docking-files/Vina-example folder has a simple docking workflow using the *dock.bash* script and the input files for the 5uf0 system, which already outputs the files in the right format for use with BAT.

The docking can also be integrated into the CELPP challenge, using the procedure from the CELPPade tutorial (https://doc.google.com/document/d/1iJcPUktbdrRftAA8cuVa32Ri1TPr2XvZVqTccDja2OMh), which the user needs to be familiarized with. The same scripts from the "internal_autodockvina_contestant" model from this tutorial can be used for docking, except for *internal_autodockvina_contestant_dock.py*. To output the docked structures suitable for use with BAT, you should use instead the *BAT_dock.py* script, which is included inside the ./docking-files/CELPP-docking folder. Once you run the CELPP docking, the necessary pdb structures for various different receptors will be inside the ./4-docking folder that was just created with the docking results.

**5.2 Co-crystal structure:** In the case of a co-crystal structure, the `calc_type`, `ligand_name` and `celpp_receptor` options in the input file have to be adjusted properly, as explained in section 4. The script *prep-crystal.tcl*, inside the ./BAT/build_files folder, is used by VMD [15] to "clean" the original file and leave only a single chain of the receptor-ligand complex. The editing of this tcl script usually not needed, but is necessary if the original pdb file has more than one chain. Keep in mind that the "MMM" identifier will be replaced by the ligand name, so it does not have to be changed beforehand.

#### 6. Adding a new protein system

BAT.py can be extended to several protein systems, by including a reference alignment file and adjusting a few parameters in the input.in file. The ./systems-library folder contains the necessary data for a few systems, and more can be requested by contacting the author directly. The user can also set up a new protein system, so below we show a few good practices, using the second BRD4 bromodomain as an example.

**6.1: Aligning the protein:** The first step is to create the reference.pdb file, so that the complex can be aligned relative to it using MUSTANG, before the BAT.py simulations are performed. If using APR, the ligand pulling direction is along the *z* axis and towards positive values (*z*+), so the ligand must have free access to the solvent along this direction. This is not needed if only DD is to be employed. The reference file is created by rotating a structure of the desired protein with a ligand bound, in this example the 5uf0 crystal structure of BRD4(2), and then saving it in the ./BAT/build_files folder as reference.pdb. One way of doing this is using VMD (http://www.ks.uiuc.edu/Training/Tutorials/vmd/vmd-tutorial.pdf), but other programs such as Chimera can also be employed for that purpose. Figure 1 shows the 5uf0 structure before (red) and after the rotation (blue), with the ligand now having access to the solvent along the *z* direction. The reference.pdb file does not need to have the same sequence as the receptor input file, so the same reference created here for for BRD4(2) can be extended to other  bromodomain homologs that share the same fold, such as BRD4(1), CREBBP and BAZ2B.

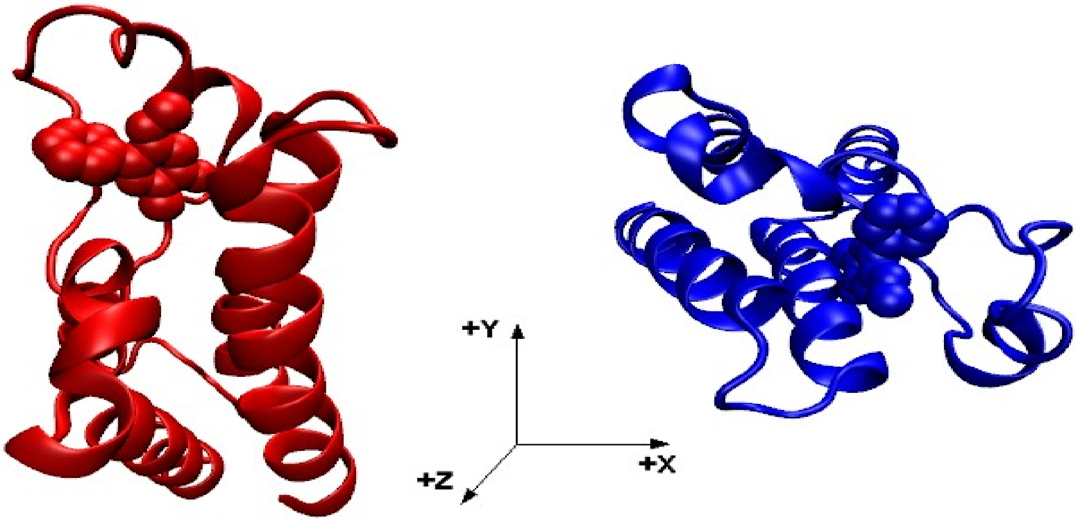


**Figure 1:** The 5uf0 structure before (red) and after (blue) the rotation, so that the latter has the ligand with free access to the solvent in the +z direction.

**6.2: Choosing the protein anchors:** Once the reference.pdb file with the desired orientation is created, it is time to choose the three protein anchor atoms. Starting from this file, choose a tentative ligand first anchor atom L1 [8], with the lowest or one of the lowest values for the *z* coordinate from all the ligand atoms, and with the *x* and *y* coordinates not too far from the center of the binding site (Figure 2). Even though the L1 anchor is going to be chosen automatically when you run BAT.py, an estimate of its location is needed to choose the protein anchors. The protein P1 anchor is then chosen using a few rules:

1 – Should be a backbone atom (CA, C or N) and part of a stable secondary structure such as an alpha-helix or a

beta sheet, always avoiding loop regions due to their increased flexibility.

2 – There should be a distance ranging from 10 Å to 15 Å between P1 and the L1 anchor atom in the *z* axis, and an absolute value between 5.0 Å and 10 Å for this same distance in the *xy* plane.

In the example for the 5uf0 structure (Figure 2), the tentative L1 atom has a {*x1 y1 z1*} distance vector from the CA atom from the protein 403 residue (P1) being equal to {-0.74 -6.16 13.03} , with the *z1* distance being 13.03 Å and the distance in the *xy* plane $x1^2 + y1^2$ being 6.20 Å. Since it satisfies our criteria, we choose :403@CA for the first protein anchor P1.

The choice of the other protein anchors P2 and P3 are chosen after the first one, and also follow a few guidelines:

1 – Should also be backbone atoms part of stable secondary structures such as alpha-helix or beta sheet, always avoiding loop regions due to their increased flexibility.

2 – The N2-P1-P2 and P1-P2-P3 angles (Figure 2) should not be close to 0 or 180 degrees (better if close to 90°), and the distance between the protein anchors should be large (at least 10 Å) . This is to avoid large forces during the simulations due to a gimbal lock. Like the ligand anchors, the positioning of the dummy atoms is done automatically, with an example shown in Figure 2. More information on their location and the full restraint setup can be found on Refs [8,9].

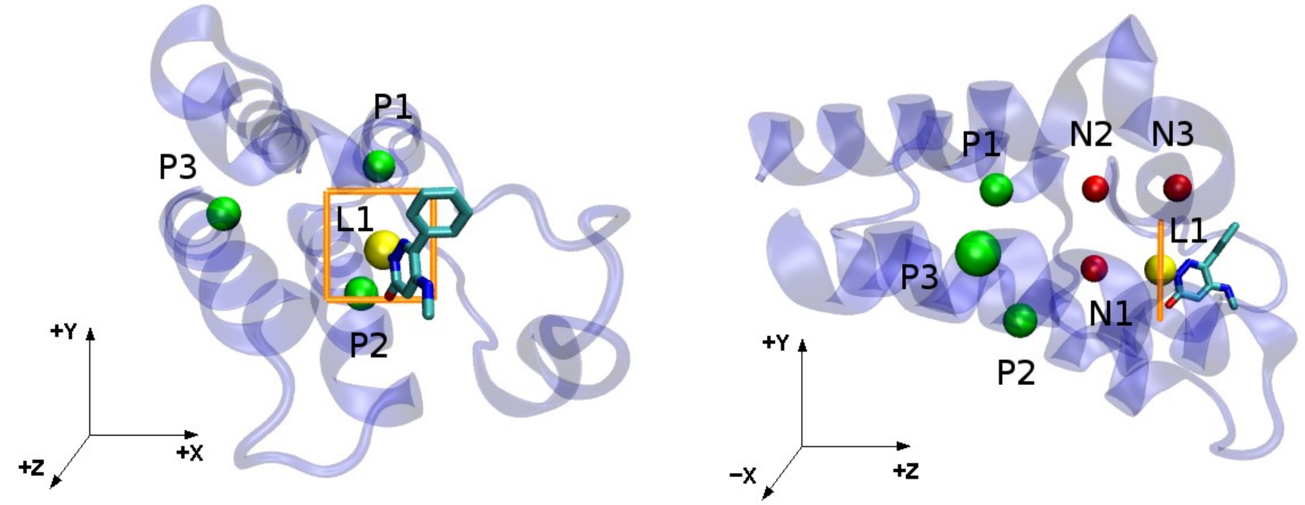


**Figure 2:** (*left*) The ligand tentative first anchor L1 (yellow) and the three protein anchors P1, P2 and P3 (green). The strike zone for L1 search is shown in orange. (*right*) The same structure, but showing the the *yz* plane. The dummy atoms N1, N2 and N3 (red), which are placed automatically when running BAT.py, are shown as an example.

**6.3: Determining the input values for ligand anchor search:** Once P1, P2 and P3 are chosen, a few more parameters are needed for the input file. The *x1* and *y1* coordinates determined above can be used for the `l1_x` and `l1_y` parameters. The minimum (`l1_z`) and maximum (`l1_zm`) P1-L1 distances in the *z* axis can have safe values of 8.50 Å and 20.0 Å, respectively. The `l1_range` parameter, which defines a “strike zone” when searching for the first ligand anchor L1, can also be safely defined as 3.0 Å (Figure 2). The `min_adis` and `max_adis` define the minimum and maximum distance between the ligand anchors, and can usually be left with values of 4.0 Å and 8.0 Å, respectively. For smaller ligands, `min_adis` might have to be reduced to 3.5 Å or even 3.0 Å, and `max_adis` could be increased in the case of larger ligands.

Even though this is a heuristic approach, if applied correctly the protein and the ligand will be properly restrained during the calculations, allowing us to obtain the absolute binding free energy of several molecules to a given protein without any further adjustments.
